# Acute buprenorphine exposure depresses neonatal respiratory chemoreflexes in the presence or absence of naloxone

**DOI:** 10.64898/2026.05.13.724975

**Authors:** Michael Frazure, Khushi Praveen, Elias Sitzmann, Emily Flanigan, Ralph Fregosi

## Abstract

Perinatal opioid exposure is a prevalent clinical concern linked to respiratory instability and adverse infant outcomes. The opioid buprenorphine is prescribed as a medication for opioid use disorder during pregnancy and used to treat neonatal opioid withdrawal syndrome, yet its direct effects on neonatal control of breathing have not been examined. Here, we asked how acute buprenorphine exposure affects breathing at rest, and during chemoreceptor stimulation. Using dual-chamber head-out plethysmography, we measured pulmonary ventilation rate (V̇_I_) and metabolic rate in awake male and female Sprague-Dawley neonatal rats on postnatal days 4-5 (P4–5) during eupnea and a hypoxic-hypercapnic (HH) challenge. The effects of buprenorphine and two opioid receptor antagonists, naloxone hydrochloride, or peripherally restricted naloxone methiodide, were assessed using a repeated measures design. V̇_I_ during eupnea and HH were markedly depressed following buprenorphine administration. Buprenorphine reduced V̇O_2_ and V̇CO_2_ and produced ventilatory equivalents for O_2_ and CO_2_ consistent with frank hypoventilation, driven by reduced breathing frequency and tidal volume (V_T_). When administered after buprenorphine, neither naloxone hydrochloride nor naloxone methiodide could rescue the buprenorphine-mediated hypoventilation in eupnea or during HH. In contrast, pre-treatment with either naloxone hydrochloride or naloxone methiodide attenuated buprenorphine-induced hypoventilation by preserving V_T_. These findings demonstrate that neonatal protective chemoreceptor reflexes are depressed by buprenorphine and suggest that pre-treatment with a peripheral opioid receptor antagonist could mitigate buprenorphine-induced hypoventilation without inducing opioid withdrawal.

**Key Points:**

- Acute buprenorphine exposure significantly depressed pulmonary ventilation rate (V̇_I_) during eupnea and hypoxic hypercapnia (HH) in awake neonatal rats.
- Buprenorphine-induced hypoventilation was driven by reduced tidal volume (V_T_) and breathing frequency.
- Buprenorphine also reduced oxygen consumption (V̇O₂) and carbon dioxide production (V̇CO₂).
- Naloxone given after buprenorphine failed to reverse hypoventilation.
- In contrast, pre-treatment with either naloxone hydrochloride or peripherally restricted naloxone methiodide mitigated buprenorphine-induced hypoventilation by preserving V_T_.

## Introduction

Increased maternal opioid use before, during, and after pregnancy has resulted in an expanding population of infants affected by perinatal opioid exposure (Hirai *et al*., 2021; Baldo, 2022). Many of these infants develop neonatal opioid withdrawal syndrome (NOWS), requiring treatment with exogenous opioids and prolonged hospitalization (Weller *et al*., 2021). Opioids are also used to control pain in critical care and post-surgical settings, which can lead to iatrogenic NOWS and further extend length of stay in the neonatal intensive care unit (Lewis *et al*., 2015). Perinatal opioid exposure is associated with numerous complications, including respiratory and feeding difficulties (Kron *et al*., 1976; Ward *et al*., 1986; Ward *et al*., 1992; LaGasse *et al*., 2003), neurodevelopmental delays (Benninger *et al*., 2020), and infant mortality (Leyenaar *et al*., 2021). Despite the seriousness of these outcomes, the direct effects of perinatal opioid exposure on neonatal control of breathing are poorly understood.

After birth, the need to regulate respiration is a critical physiological transition for the newborn (Whitaker-Fornek & Levitt, 2025). Blood gas homeostasis requires dynamic modulation of ventilation to meet fluctuating metabolic demands (Chowdhuri & Badr, 2017). This process is mediated by central and peripheral chemoreflex pathways that provide excitatory input to brainstem respiratory centers controlling the rate and depth of breathing (Krohn *et al*., 2023).

Central chemoreceptors in the ventrolateral medulla sense pH changes in the cerebrospinal fluid arising from alterations in the arterial partial pressure of carbon dioxide (PaCO₂) (Nattie & Li, 2012; Guyenet & Bayliss, 2022). Peripheral chemoreceptors within the carotid bodies and aortic arch respond to changes in the arterial partial pressure of oxygen (PaO₂) and transmit ascending information to brainstem respiratory centers via the glossopharyngeal and vagus nerves, respectively (Prabhakar & Peng, 2004; Frazure *et al*., 2026). Although activation of either chemoreceptor pathway alone is sufficient to increase ventilation, conditions that produce hypoventilation (e.g., prolonged apnea, airway obstruction) rarely produce hypoxia or hypercapnia in isolation (Lanphier & Rahn, 1963; Levitzky, 2008; Smith *et al*., 2010). Accordingly, here we study the ventilatory response to hypoxic hypercapnia (HH), which simultaneously activates central and peripheral chemoreceptors (Boyd *et al*., 2022).

Available evidence indicates that the developing respiratory control network is sensitive to opioids. Prenatal methadone exposure increases apnea frequency and duration and alters hypoxic ventilatory control in neonatal rats (Hocker *et al*., 2021), while acute methadone administration depresses respiratory frequency *in vivo* and abolishes respiratory activity in isolated brainstem-spinal cord preparations (Beyeler *et al*., 2023). Prenatal morphine exposure blunts the hypoxic and hypercapnic ventilatory responses in neonatal rats (Osborne *et al*., 2022), and acute morphine produces prolonged respiratory pauses, metabolic depression, and impaired thermoregulation (Kesavan *et al*., 2014). Together, these data indicate that opioids used clinically disrupt multiple aspects of neonatal breathing, including rhythm generation and chemoreflex responses. However, important gaps remain in our understanding of how specific opioids, including buprenorphine, affect neonatal breathing.

Buprenorphine is a semisynthetic, partial μ-opioid receptor agonist thought to have a favorable safety profile, including a ceiling effect on respiratory depression (Walsh *et al*., 1994; Dahan *et al*., 2005; Gudin & Fudin, 2020). In recent years, buprenorphine has been increasingly prescribed as a medication for opioid use disorder (MOUD) during pregnancy and as a treatment for NOWS (Nguemeni Tiako *et al*., 2022; Suarez *et al*., 2022). While several studies suggest that buprenorphine exposure negatively affects neurodevelopment in general (Schlagal *et al*., 2021; Elam *et al*., 2022; Nieto-Estevez *et al*., 2022; Nyberg *et al*., 2025a; Nyberg *et al*., 2025b), its effects on neonatal respiratory control have not been examined. Given the prevalence of perinatal buprenorphine exposure, there is a critical need to understand how this specific opioid affects the control of breathing during early development. For this reason, we measured the effects of buprenorphine, in the presence or absence of opioid receptor antagonists, on ventilation, metabolic rate and chemoreflex function in neonatal rats.

## Methods

### Ethical Approval

All procedures were approved by the University of Arizona Animal Care and Use Committee (Protocol # 2024-1184) and were performed in accordance with the American Physiological Society’s Animal Care Guidelines, the National Institutes of Health (NIH) guide for the use and care of laboratory animals, ARRIVE (Animal Research: Reporting of *In Vivo* Experiments) guidelines and the stance of *The Journal of Physiology* (O’Halloran, 2024).

### Experimental Animals

We used a total of 41 neonatal Sprague-Dawley rats of both sexes (22 male) aged postnatal day 4-5 (P4-5), with an average weight of 11.8 ± 1.9 g. The neonates were derived from nine outbred Sprague-Dawley timed-pregnant dams obtained from Charles River Laboratories (Wilmington, MA, USA). Two to six pups, half male and half female, were taken from each litter. Neonates were housed with their mothers and litter mates under a 12:12 h light-dark cycle with food and water available *ad libitum*. Upon completion of *in vivo* plethysmography, animals were deeply anaesthetized on ice (confirmed by absence of withdrawal reflexes) and decapitated, in line with approved euthanasia procedures.

### Drugs

Buprenorphine hydrochloride was obtained from United States Pharmacopeia (Bethesda, MD, USA); naloxone hydrochloride from Hello Bio Inc. (Princeton, NJ, USA); and naloxone methiodide from Santa Cruz Biotechnology (Dallas, TX, USA). All drugs were prepared in deionized water (dH₂O); naloxone hydrochloride and naloxone methiodide readily dissolved, and buprenorphine was brought into solution with the addition of a single drop of hydrochloric acid. Stock solutions were prepared at concentrations of 1 mg/mL for buprenorphine, 2 mg/mL for naloxone hydrochloride, and 4 mg/mL for naloxone methiodide, aliquoted, and stored at −20 °C until the day of each experiment. Drugs were administered subcutaneously at doses of 0.3 mg/kg (buprenorphine), 0.6 mg/kg (naloxone hydrochloride), and 1 mg/kg (naloxone methiodide) using a Hamilton syringe, with injection volumes ≤ 5 μL. In time and sham-controls, saline (0.9% NaCl in dH₂O) was administered subcutaneously in volumes matched to drug injections to account for potential effects of injection and time.

### In vivo plethysmography

We measured ventilatory and metabolic parameters in awake neonatal rats using a custom dual-chamber head-out plethysmography system (depicted in Fig. 1), which has been validated and described previously (Boyd *et al*., 2022). An inner, head-out plethysmograph was connected to a pneumotachometer, which in turn was connected to the positive and negative ports of a pressure transducer (± 2 cm H₂O range; Validyne DP45-16, Northridge, CA) and amplified using a Validyne carrier demodulator. The pressure signal, proportional to airflow, was routed in parallel to an analog-to-digital (A/D) board (Cambridge Electronic Design). The system was calibrated using known flows and volumes, and the inspiratory component of the airflow signal was rectified and integrated to derive inspired tidal volume (V_T_). A gas analyzer (iWorx GA-200) continuously pulled air through the outer chamber and a Dri-Rite canister at a rate of 150 mL/min, and sampled the concentrations of O_2_ and CO_2_ in the effluent gas. The recording chamber was positioned on a heating pad which was servo-controlled by monitoring the internal chamber temperature with a thermocouple connected to a far infra-red controller (Kent Scientific). Chamber temperature was maintained at 31-33 °C, corresponding to normal nesting conditions for neonatal rats (Mortola, 1984).

**Figure 1.**
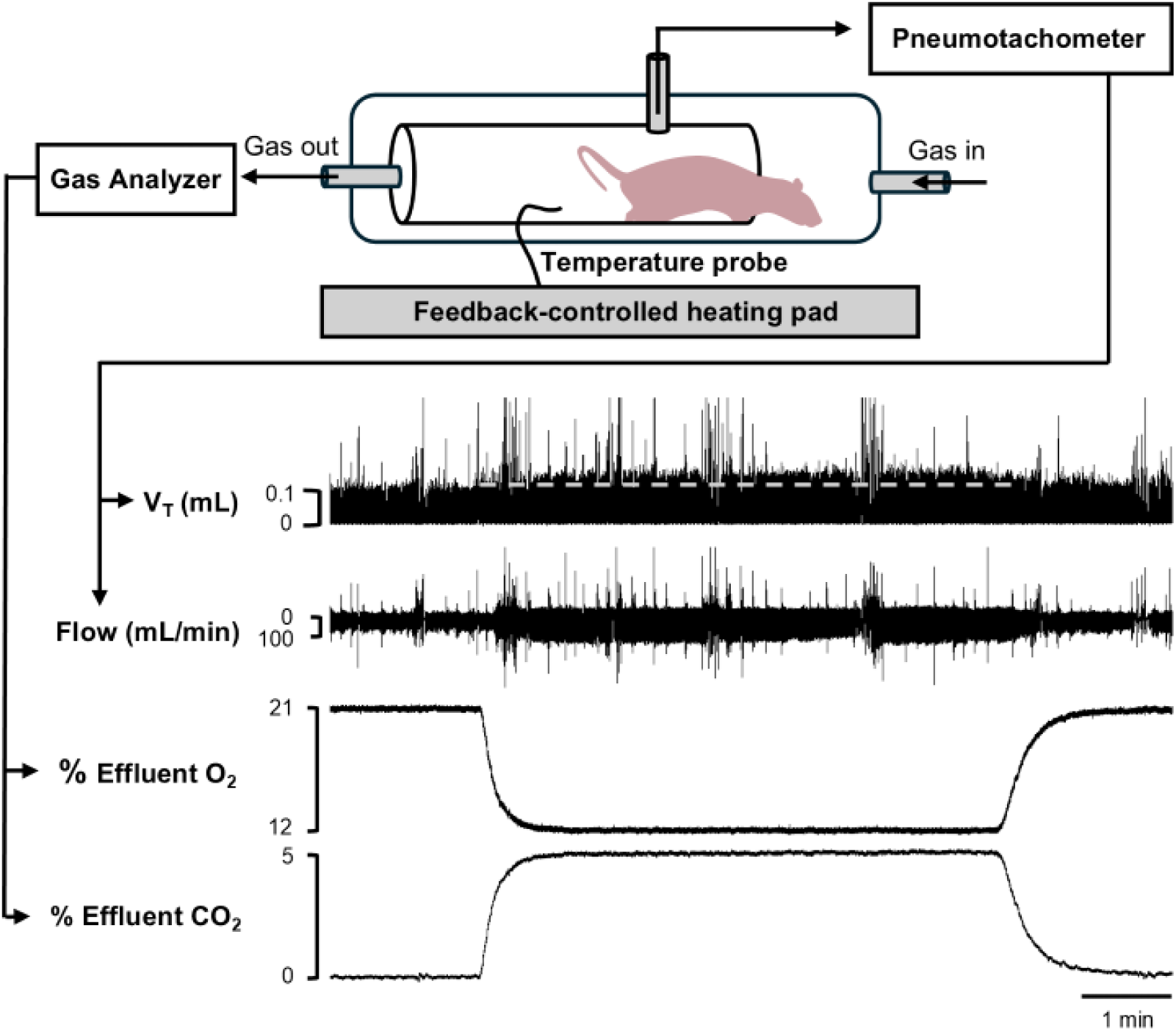
Dual chamber head-out plethysmography system for measuring ventilatory and metabolic parameters in neonatal rats. The schematic depicts a rat pup in a head-out plethysmography chamber nested within an outer chamber. This configuration permits manipulation of the inspired gas mixture while enabling measurement of pressure changes produced by thoracic expansion. Either room air (RA; 21% O_2_, 0% O_2_) or a hypoxic-hypercapnic (HH) gas mixture (12% O_2_, 5% CO_2_) enters the system at a constant flow rate. Breathing-related pressure changes within the inner chamber are detected by a pneumotachometer and pressure transducer, allowing calculation of respiratory frequency, tidal volume (V_T_), and pulmonary ventilation rate (V̇_I_). Effluent gas is sampled by a gas analyzer; changes in the percent O_2_ and CO_2_ in the chamber effluent, relative to the inspired values, are used to calculate oxygen consumption (V̇O_2_) and carbon dioxide production (V̇CO_2_). Representative traces show airflow and integrated V_T_ during RA breathing and a five-minute HH challenge followed by a recovery. The dashed white line highlights the increase in V_T_ during the HH challenge, whereas very large amplitude signals indicate movement artifact.

Following calibration, pups were placed into the head-out plethysmograph and a seal formed around the neck using a thin nitrile diaphragm and vacuum grease. The plethysmograph containing the pup was then placed into the larger outer chamber. After confirming a stable airflow signal, animals were acclimated to the chamber for 15 min prior to data collection. A 5-min period of eupneic room air (RA) breathing was followed by a 5-min hypoxic–hypercapnic (HH) challenge using a 12% O₂/5% CO₂ gas mixture delivered via a Matheson rotameter at 155 mL/min. At this inflow rate, chamber gas concentrations reached 12% O₂ and 5% CO₂ within approximately 1 min and were maintained throughout the 5-min challenge (Fig. 1). After the HH challenge, the chamber refilled with RA within 1 min, followed by a 5-min recovery period. To test the effects of buprenorphine in the presence and absence of opioid receptor antagonists, three primary experimental protocols were performed.

### Experiment I: Buprenorphine followed by naloxone hydrochloride

To assess the effects of buprenorphine alone, and to determine whether these effects could be reversed by a competitive opioid receptor antagonist, experiments were first performed in *n* = 10 neonatal Sprague Dawley rats (6 males; P4-5). Control measurements were obtained during RA breathing and the HH challenge. After recovery in RA, buprenorphine was administered, and measurements were repeated 30 min later. Naloxone hydrochloride was subsequently administered, and measurements repeated after an additional 30 min. Here and throughout, injections were administered into the scruff of the neck by briefly removing the outer chamber cap. After checking and adjusting the neck-seal as needed, the outer chamber cap was replaced and recording resumed.

### Experiment II: Naloxone hydrochloride followed by buprenorphine

Next, experiments were performed in *n* =10 neonatal Sprague Dawley rats (5 males; P4-5) to assess the effects of naloxone hydrochloride alone, and to determine whether pre-treatment with a competitive antagonist could mitigate the respiratory depressant effects of buprenorphine. Control measurements were obtained during RA breathing and the HH challenge. Following recovery in RA, naloxone hydrochloride was administered, and measurements were repeated 30 min later. Buprenorphine was subsequently administered, and measurements were repeated after an additional 30 min.

### Experiment III: Naloxone methiodide followed by buprenorphine

Finally, experiments were performed in *n* = 8 neonatal Sprague Dawley rats (4 males; P4-5) to determine whether pre-treatment with a peripherally restricted opioid receptor antagonist could mitigate the respiratory depressant effects of buprenorphine. Control measurements were obtained during RA breathing and the HH challenge. Following recovery in RA, naloxone methiodide was administered, and measurements were repeated 30 min later. Buprenorphine was subsequently administered, and measurements were repeated after an additional 30 min.

### Supplemental experiments

Experiments were also performed in *n* = 7 neonatal Sprague Dawley rats (4 males; P4-5) in which buprenorphine administration was followed by peripherally restricted naloxone methiodide to control for the order of drug administration in Experiment III. These results are presented in the Supplemental Materials (Supplemental Fig. 1, Supplemental Table 1). Time and sham-controls were conducted in *n* = 6 neonatal Sprague Dawley rats (3 males; P4-5) using two saline injections instead of drug injections, with measurements obtained over the same time course as Experiments I–III (Supplemental Fig. 2).

### Statistical analysis

The primary outcome measures made in all experiments were pulmonary ventilation rate (V̇_I_), tidal volume (V_T_), breathing frequency, inspiratory and expiratory phase durations (T_I_ and T_E_, respectively), apnea frequency and duration, oxygen consumption (V̇O₂), carbon dioxide production (V̇CO₂), the ventilatory equivalent for oxygen (V̇_I_/V̇O₂) and the ventilatory equivalent for carbon dioxide (V̇_I_/V̇CO₂). Apneas were defined as inter-breath intervals > 2× the mean T_E_ for a given condition. V_T_, breathing frequency, T_I_, T_E_, and apnea frequency and duration were measured using Spike2 software. V̇_I_ was calculated offline as the product of V_T_ and breathing frequency, normalized to bodyweight. Values for V_T_ and V̇_I_ are reported under ATPS conditions. Oxygen consumption (V̇O₂) and carbon dioxide production (V̇CO₂) were calculated as:

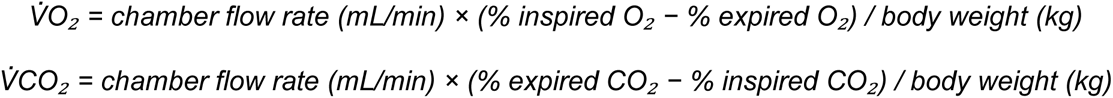

Data were entered into Excel spreadsheets (Microsoft Corp., Redmond, WA, USA) and exported to Prism for statistical analysis (GraphPad Software, San Diego, CA, USA). Sex differences were assessed using a three way repeated measures ANOVA with biological sex as the between-subjects factor, and inspired gas mixture and drug treatment as within-subjects factors, as each animal was studied across gas and drug conditions and served as its own control. This analysis revealed no significant effect of sex (Supplemental Fig. 3); therefore, data from male and female animals were pooled for analyses presented in the main manuscript.

V̇_I_, V_T_, breathing frequency, T_I_, T_E_, and apnea frequency and duration were analyzed using two-way repeated measures ANOVA, with gas mixture and drug treatment as within-subject factors, as each animal was studied across gas and drug conditions and served as its own control. While the mean ± SD values for each minute of the HH challenge are shown (Figs. 2, 5, 8), statistical analyses focused on comparisons between eupneic RA breathing and the first minute of the HH challenge at the peak of the hypoxic-hypercapnic ventilatory response (HHVR).

**Figure 2.**
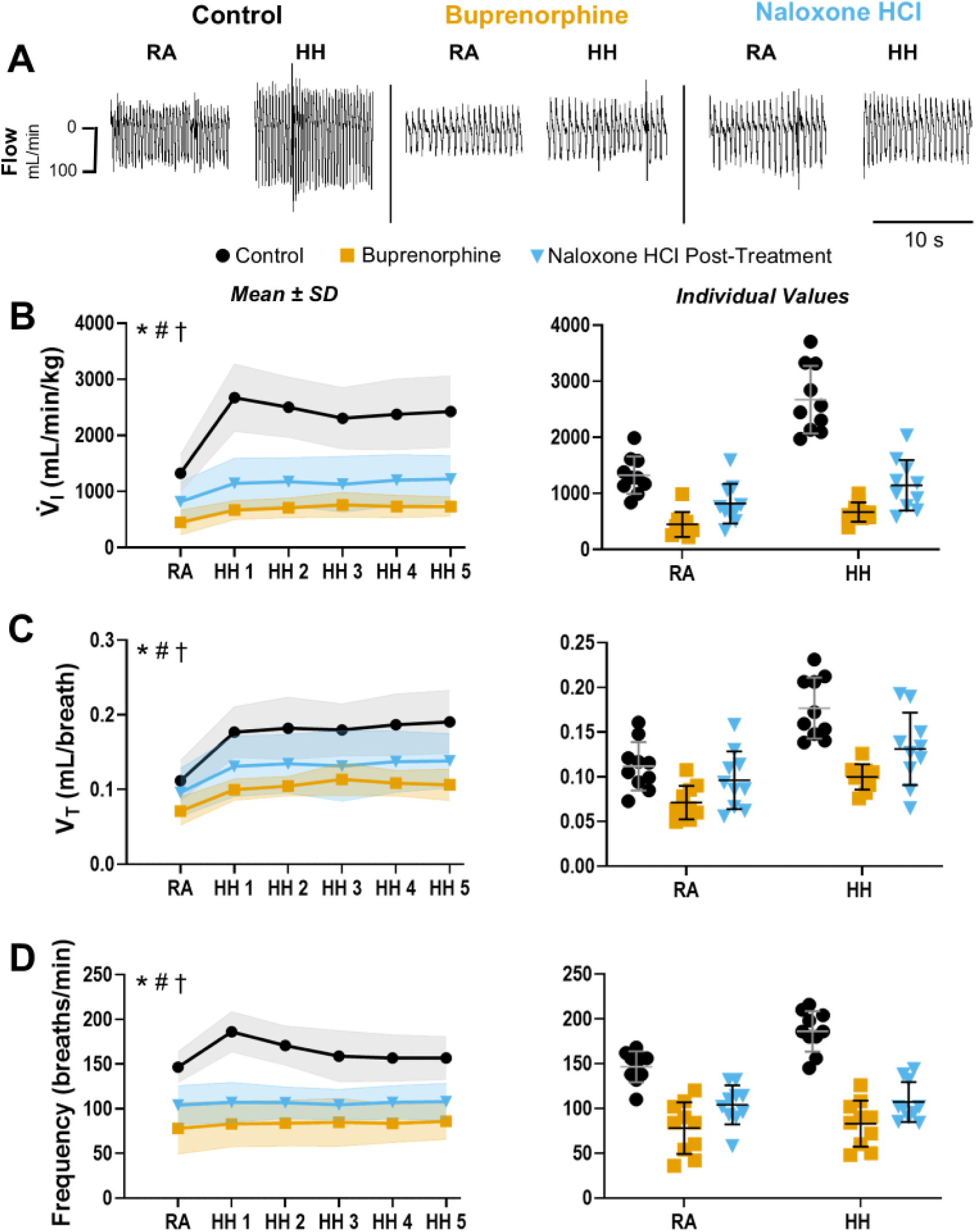
Acute buprenorphine exposure depresses the hypoxic-hypercapnic ventilatory response in neonatal rats. V̇_I_, V_T_ and breathing frequency were measured using dual-chamber plethysmography in neonatal Sprague Dawley rats (*n*=10, 6 male; P4-5) during RA breathing and a 5-min HH challenge (HH1-5). Measurements were obtained before and after administration of buprenorphine, followed by naloxone hydrochloride (Naloxone HCl). **Panel A:** Representative plethysmography traces illustrate the effects of gas mixture (RA, HH) and drug condition (Control, Buprenorphine, Naloxone HCl) on breathing. **Panels B–D:** Group data (left) and individual replicates (right) show the effects of drug and gas mixture on V̇_I_ (B), V_T_ (C), and breathing frequency (D). Under control conditions, there was a robust hypoxic hypercapnic ventilatory response (HHVR). Buprenorphine reduced resting ventilation and markedly attenuated the HHVR, blunting the V_T_ response and abolishing the frequency response. Naloxone hydrochloride produced modest increases in V_T_ and frequency relative to buprenorphine alone but failed to restore V̇_I_ to baseline. Data are presented as mean ± SD. Statistical analyses were performed using two-way repeated-measures ANOVA with Tukey’s HSD post-hoc tests and statistical significance defined as *p* < 0.05. Significant ANOVA effects are indicated in the upper left of each panel: Gas (*); Drug (#); Gas × drug interaction (†). Detailed statistics are reported in the Results, and post-hoc comparisons are listed in Table 1.

**Table 1.**
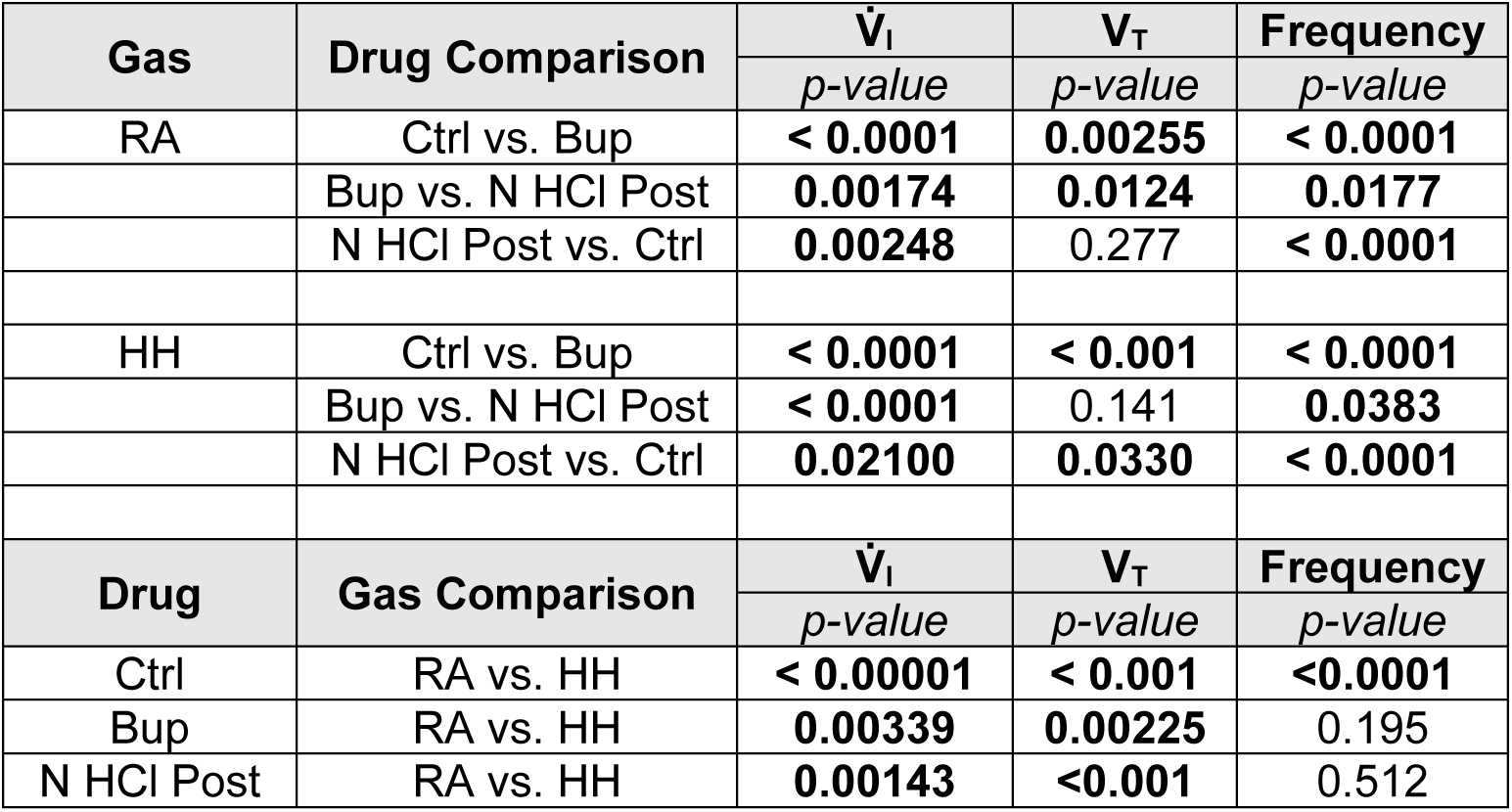
Post-hoc comparisons of respiratory parameters across drug treatments and gas mixtures in Experiment I. *P*-values shown in bold when significant (*p* < 0.05). *RA = Room air; HH = Hypoxic-hypercapnic challenge; Ctrl = Control; Bup = Buprenorphine; N HCl Post = Naloxone hydrochloride post-treatment*.

**Table 2.** Post-hoc comparisons of respiratory parameters across drug treatments and gas mixtures in Experiment II. *P*-values shown in bold when significant (*p* < 0.05). *RA = Room air; HH = Hypoxic-hypercapnic challenge; Ctrl = Control; N HCl Pre = Naloxone hydrochloride pre-treatment; Bup = Buprenorphine*.

| Gas | Drug Comparison | $\dot{V}_I$ | $V_T$ | Frequency |
| --- | --- | --- | --- | --- |
|  |  | *p*-value | *p*-value | *p*-value |
| RA | Ctrl vs. N HCl Pre | 0.622 | 0.089 | 0.273 |
|  | N HCl Pre vs. Bup | **0.00453** | **0.00484** | **0.00790** |
|  | Bup vs. Ctrl | **0.0107** | 0.144 | **0.00352** |
| HH | Ctrl vs. N HCl Pre | 0.0501 | 0.671 | 0.187 |
|  | N HCl Pre vs. Bup | **0.00233** | 0.378 | **0.00187** |
|  | Bup vs. Ctrl | **0.00113** | 0.323 | **< 0.0001** |
| Drug | Gas Comparison | $\dot{V}_I$ | $V_T$ | Frequency |
|  |  | *p*-value | *p*-value | *p*-value |
| Ctrl | RA vs. HH | **< 0.0001** | **< 0.0001** | **<0.0001** |
| N HCl Pre | RA vs. HH | **< 0.0001** | **0.00205** | **<0.001** |
| Bup | RA vs. HH | **0.00111** | **<0.001** | **0.0201** |

**Table 3.**
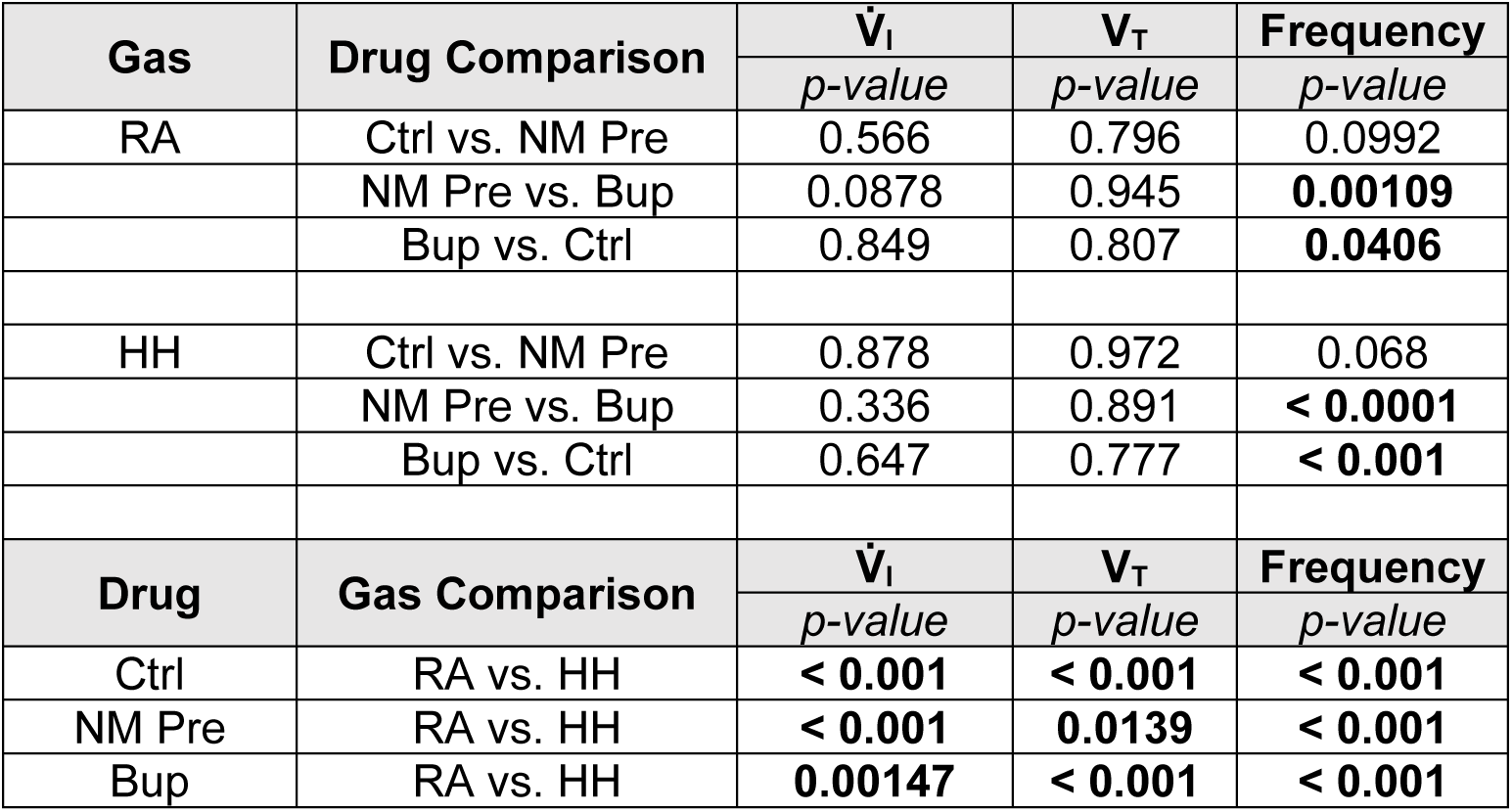
Post-hoc comparisons of respiratory parameters across drug treatments and gas mixtures in Experiment III. *P*-values shown in bold when significant (*p* < 0.05). *RA = Room air; HH = Hypoxic-hypercapnic challenge; Ctrl = Control; NM Pre = Naloxone methiodide pre-treatment; Bup = Buprenorphine*.

| Gas | Drug Comparison | $\dot{V}_I$ | $V_T$ | Frequency |
| --- | --- | --- | --- | --- |
|  |  | *p*-value | *p*-value | *p*-value |
| RA | Ctrl vs. NM Pre | 0.566 | 0.796 | 0.0992 |
|  | NM Pre vs. Bup | 0.0878 | 0.945 | **0.00109** |
|  | Bup vs. Ctrl | 0.849 | 0.807 | **0.0406** |
| HH | Ctrl vs. NM Pre | 0.878 | 0.972 | 0.068 |
|  | NM Pre vs. Bup | 0.336 | 0.891 | **< 0.0001** |
|  | Bup vs. Ctrl | 0.647 | 0.777 | **< 0.001** |
| Drug | Gas Comparison | $\dot{V}_I$ | $V_T$ | Frequency |
|  |  | *p*-value | *p*-value | *p*-value |
| Ctrl | RA vs. HH | **< 0.001** | **< 0.001** | **< 0.001** |
| NM Pre | RA vs. HH | **< 0.001** | **0.0139** | **< 0.001** |
| Bup | RA vs. HH | **0.00147** | **< 0.001** | **< 0.001** |

V̇O₂, V̇CO₂, V̇_I_/V̇O₂, and V̇_I_/V̇CO₂ were calculated during steady-state RA breathing and analyzed using one-way repeated measures ANOVA, with drug treatment as the within-subjects factor; each animal was studied across drug conditions and served as its own control. Throughout, when significant ANOVA effects or interactions were detected, post hoc comparisons were performed using Tukey’s honestly significant difference (HSD) test corrected for multiple comparisons. For all analyses, statistical significance was defined as *p* < 0.05. Data presented as mean ± SD.

## Results

### Experiment I: Buprenorphine followed by naloxone hydrochloride

#### V̇_I_, V_T_ and breathing frequency

Two-way repeated measures ANOVA revealed significant effects of gas mixture (F_1,9_ = 103.0, *p* < 0.0001), drug (F_2,18_ = 86.98, *p* < 0.0001), and a gas × drug interaction (F_2,18_ = 37.08, *p* < 0.0001) for V̇_I_. Significant effects of gas mixture (F_1,9_= 153.8, *p* < 0.0001), drug (F_2,18_ = 16.54, *p* < 0.001), and a gas × drug interaction F_2,18_ = 5.51, *p* = 0.0165) were also observed for V_T_. There were also significant effects of gas mixture (F_1,9_ = 44.10, *p* < 0.0001), drug (F_2,18_ = 86.82 p < 0.0001), and a gas × treatment interaction (F_2,18_ = 24.61, *p* < 0.0001) for breathing frequency. Data are shown in Figure 2, with post-hoc comparisons listed in Table 1.

Buprenorphine significantly reduced V̇_I_ during RA breathing via combined reductions in V_T_ and respiratory frequency (Fig. 2). Under control conditions, HH elicited a robust increase in V̇_I_ driven by elevations in both V_T_ and breathing frequency. Following buprenorphine administration, the HHVR was markedly attenuated due to combined depression of V_T_ and breathing frequency. Notably, the frequency response to HH was abolished (Fig. 2D), whereas V_T_ still exhibited a small but significant increase (Fig. 2C). Subsequent naloxone hydrochloride did not restore V̇_I_ during eupnea or HH to baseline, but did produce subtle effects on V_T_ and breathing frequency relative to buprenorphine (Fig. 2C and 2D). During RA breathing, naloxone hydrochloride increased V_T_ toward baseline; however, the ability to increase V_T_ in response to HH was diminished. Naloxone hydrochloride increased breathing frequency during both RA and HH, although it remained significantly lower than control values.

#### Respiratory pattern

Two-way repeated measures ANOVA revealed significant effects of drug (F_2,18_ = 14.91, *p* < 0.001) on T_I_, which increased following buprenorphine and remained above control following naloxone hydrochloride (Fig. 3B). Significant effects of both gas mixture (F_1,9_ = 6.72, *p* = 0.0291) and drug (F_2,18_ = 8.89, *p* = 0.0138) were observed for T_E_. Under control conditions, the frequency response to HH was driven by shortened T_E_. Buprenorphine prolonged T_E_ during RA breathing, an effect that persisted throughout the HH challenge and abolished the normal frequency response (Fig. 3C). Naloxone hydrochloride did not restore T_I_ or T_E_ to baseline levels. Two-way repeated-measures ANOVA revealed no significant effects of gas mixture or drug on apnea frequency or duration. Under control conditions, apnea frequency during RA breathing averaged 0.7 ± 0.4 apneas per minute, with a mean duration of 0.5 ± 0.3 s.

**Figure 3.**
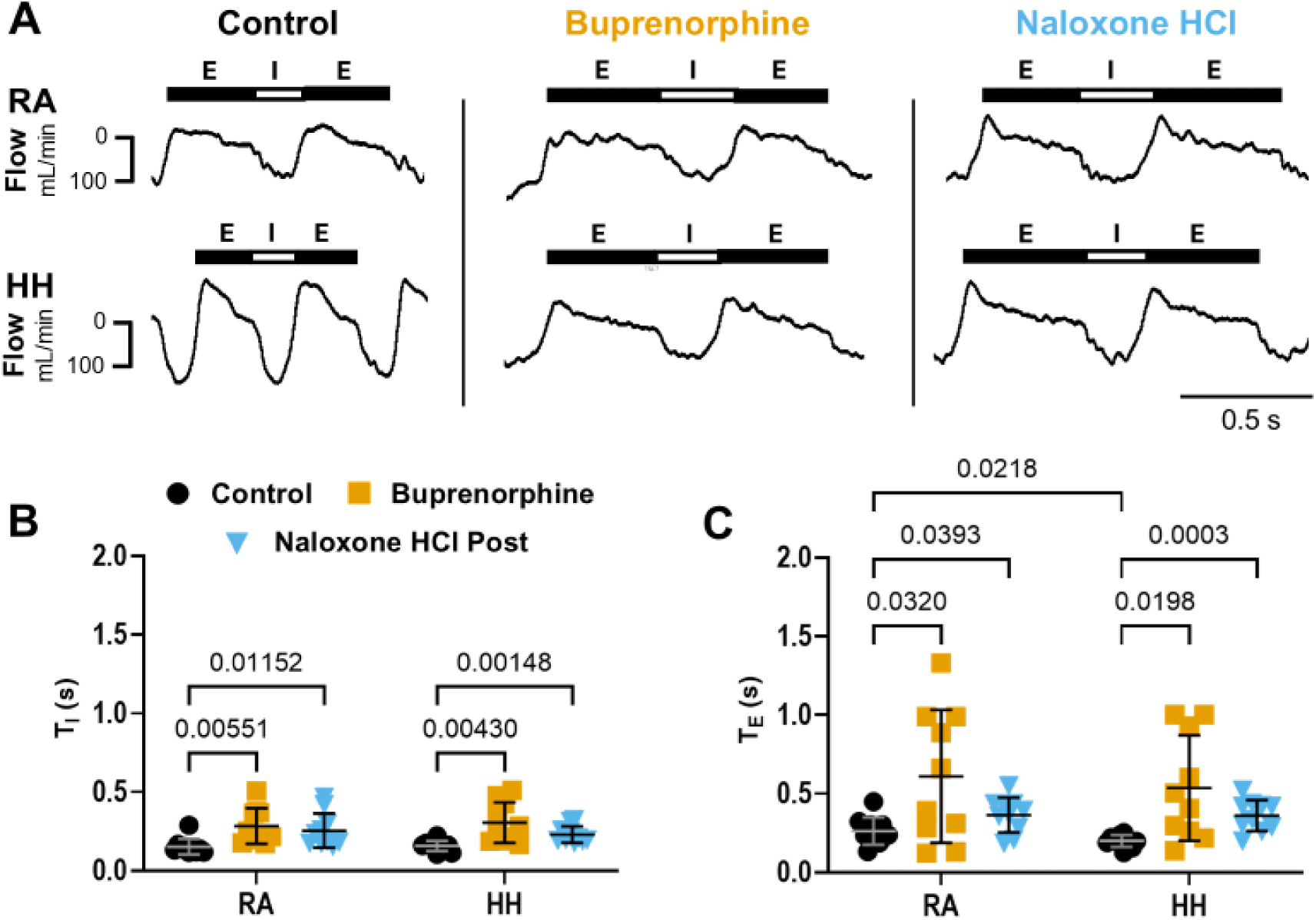
Buprenorphine-induced breathing frequency depression is characterized by prolonged T_I_ and T_E_. T_I_ and T_E_ were measured using dual-chamber plethysmography in neonatal Sprague Dawley rats (*n*=10, 6 male; P4-5) during RA breathing and a HH challenge. Measurements were obtained before and after buprenorphine administration, followed by naloxone hydrochloride (Naloxone HCl). **Panel A:** Representative plethysmography traces illustrate breathing patterns across drug conditions (Control, Buprenorphine, Naloxone HCl) and gas mixtures (RA, HH). Troughs and plateaus represent inspiration (I) and expiration (E), respectively. **Panels B and C:** Plots show group data for T_I_ (B) and T_E_ (C). T_I_ increased following buprenorphine and remained above the control duration after naloxone. Under control conditions, T_E_ shortens in response to HH. Buprenorphine prolonged T_E_ and eliminated the frequency response to HH, and naloxone hydrochloride did not reverse these changes. Data are presented as mean ± SD. Statistical analyses were performed using two-way repeated-measures ANOVA with Tukey’s HSD post-hoc tests and statistical significance defined as *p* < 0.05. Significant post-hoc comparisons are indicated on the plots.

#### Metabolic rate

One-way repeated measures ANOVA revealed a significant effect of drug on V̇O₂ (F_2,18_ = 13.17, *p* < 0.001), V̇CO₂ (F_2,18_ = 10.78, *p* = 0.00133), V̇_I_/V̇O₂ (F_2,18_ = 11.81, *p* < 0.001), and V̇_I_/V̇CO₂ (F_2,18_ = 37.41, *p* < 0.0001). Buprenorphine administration reduced V̇O₂ and V̇CO₂, and the ventilatory equivalents for O_2_ and CO_2_ (V̇I/V̇O₂ and V̇I/V̇CO₂). These data confirm that the decrease in V̇_I_ was driven by buprenorphine’s effect on the drive to breathe, rather than a decline in metabolic rate. Although subsequent naloxone administration increased V̇O₂ and V̇CO₂, V̇I/V̇O₂ and V̇I/V̇CO₂ remained significantly reduced, as V̇_I_ failed to recover. Data and significant post hoc comparisons are shown in Figure 4.

**Figure 4.**
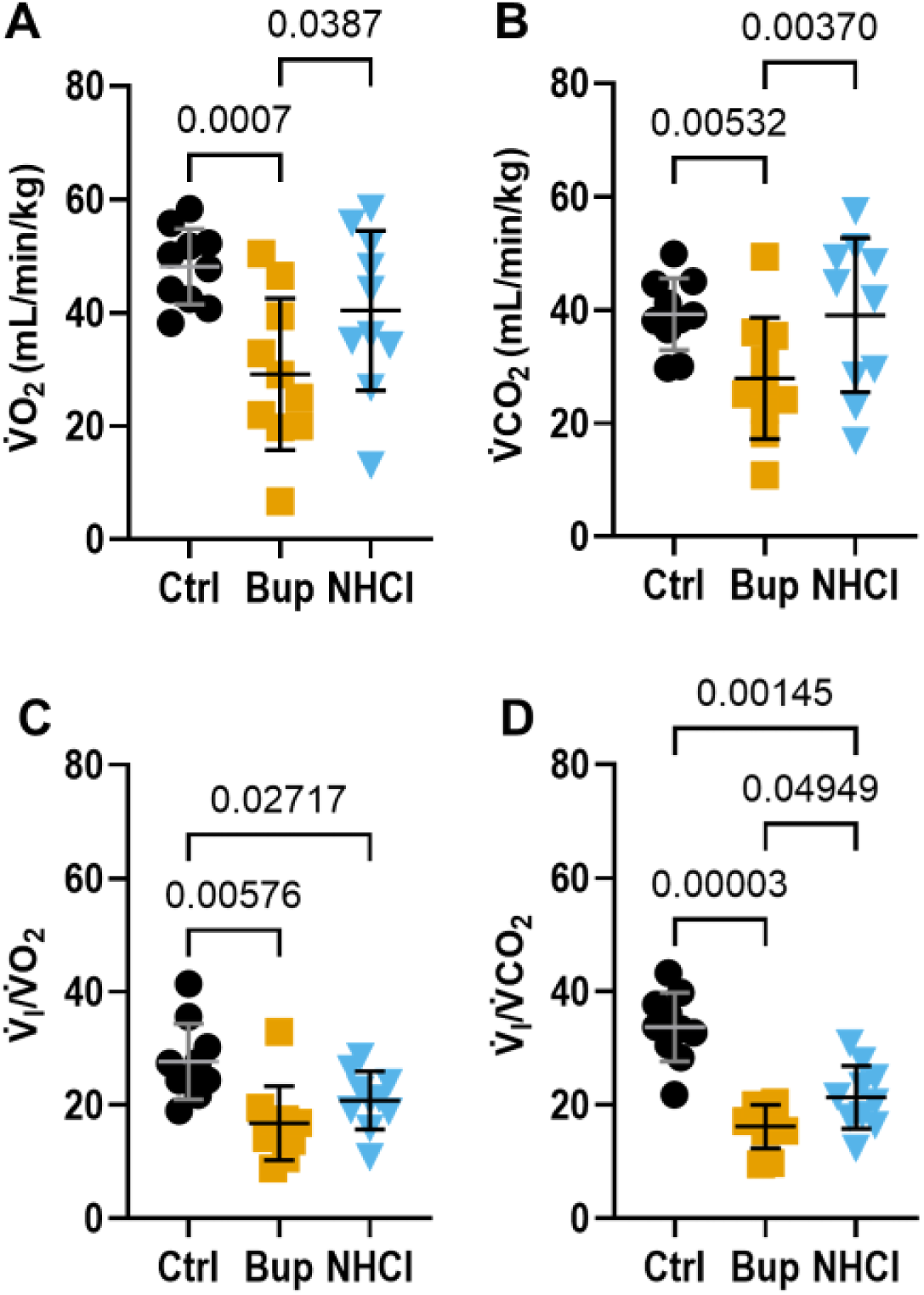
Acute buprenorphine depresses metabolic rate and ventilatory equivalents for O_2_ and CO_2_. V̇O₂, V̇CO₂, V̇I/V̇O₂ and V̇I/V̇CO₂ were measured using dual-chamber plethysmography in neonatal Sprague Dawley rats (*n*=10, 6 male; P4-5) during RA breathing. Measurements were obtained under control conditions (Ctrl) and repeated following administration of buprenorphine (Bup), followed by naloxone hydrochloride (NHCl). **Panels A-D:** Group data showing V̇_I_ (A), V̇CO₂ (B), V̇I/V̇O₂ (C) and V̇I/V̇CO₂ (D) across drug conditions. Buprenorphine significantly reduced V̇O₂ and V̇CO₂, indicating depression of metabolic rate. V̇I/V̇O₂ and V̇I/V̇CO₂ were also reduced following buprenorphine, indicating hypoventilation. Despite restoration of V̇O₂ and V̇CO₂ following naloxone, V̇I/V̇O₂ and V̇I/V̇CO₂ remained reduced. Data are presented as mean ± SD. Statistical analyses were performed using one-way repeated-measures ANOVA with Tukey’s HSD post-hoc tests and statistical significance defined as *p* < 0.05. Significant post-hoc comparisons are indicated on the plots.

### Experiment II: Naloxone hydrochloride followed by buprenorphine

#### V̇_I_, V_T_ and breathing frequency

Two-way repeated measures ANOVA revealed significant effects of gas mixture (F_1,9_ = 53.32, *p* < 0.0001), drug (F_2,18_ = 19.65, *p* < 0.001), and a gas × drug interaction (F_2,18_ = 21.17, *p* < 0.001) for V̇_I_. A significant effect of gas mixture (F_1,9_ = 50.71, *p* < 0.0001) was observed on V_T_. Significant effects of gas mixture (F_1,9_ = 54.24, *p* < 0.0001), drug (F_2,18_ = 28, p < 0.0001), and a gas × drug interaction (F_2,18_ = 16.48, p < 0.001) were observed for breathing frequency. Data are shown in Figure 5, with post-hoc comparisons listed in Table 2.

**Figure 5.**
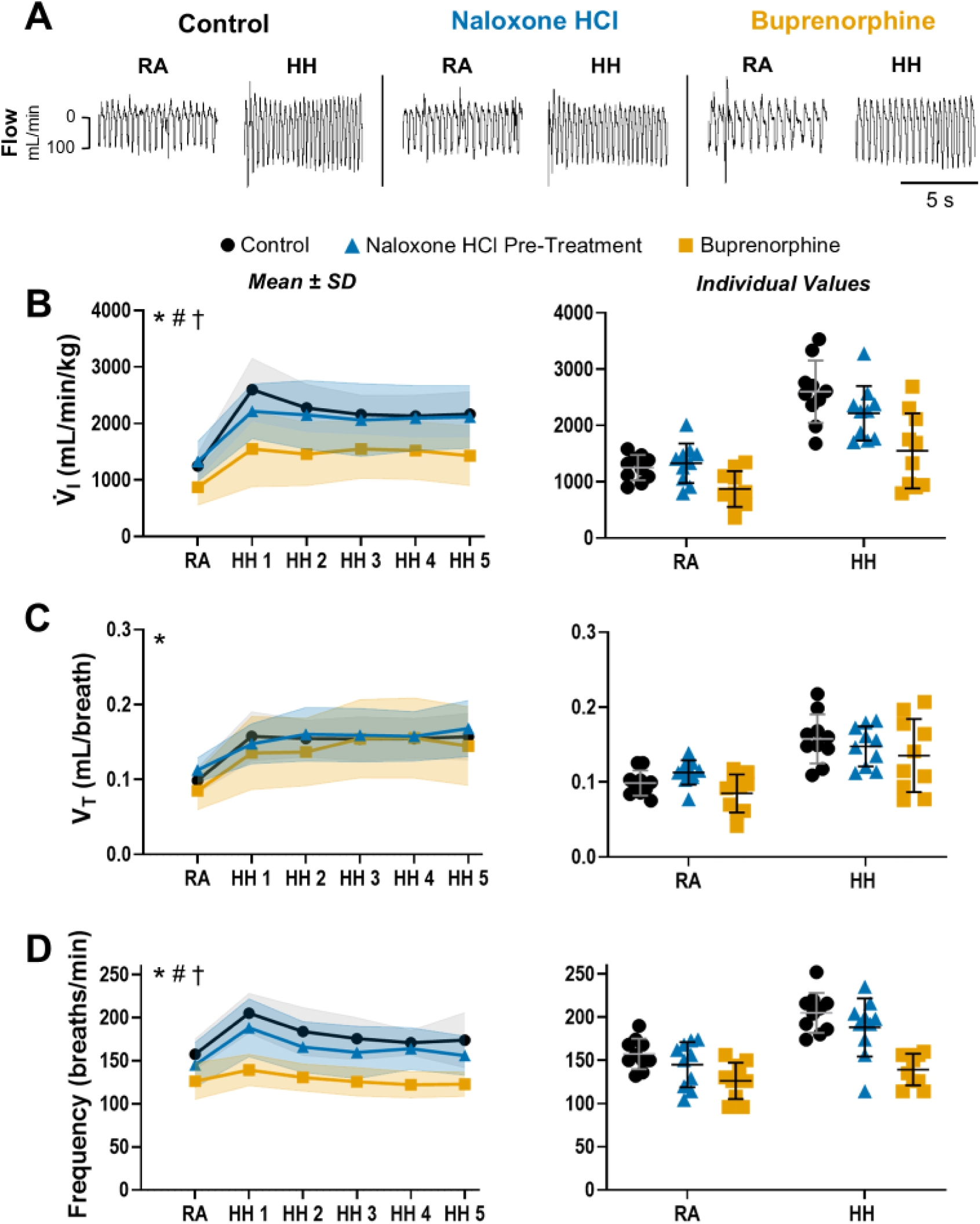
Naloxone hydrochloride pre-treatment attenuates buprenorphine-induced hypoventilation. V̇_I_, V_T_ and breathing frequency were measured using dual-chamber plethysmography in neonatal Sprague Dawley rats (*n*=10, 5 male; P4-5) during RA breathing and a 5-min HH challenge (HH1-5). Measurements were obtained before and after administration of naloxone hydrochloride (Naloxone HCl), followed by buprenorphine. **Panel A:** Representative plethysmography traces illustrate the effects of gas mixture (RA, HH) and drug condition (Control, Naloxone HCl, Buprenorphine) on breathing. **Panels B–D:** Group data (left) with individual replicates (right) show the effects of drug and gas mixture on V̇_I_ (B), V_T_ (C), and breathing frequency (D). Naloxone alone did not affect V̇_I_, V_T_, or breathing frequency during RA or HH. Buprenorphine subsequently reduced V̇_I_ at rest and during HH via frequency depression, whereas naloxone pre-treatment prevented V_T_ depression. Data are presented as mean ± SD. Statistical analyses were performed using two-way repeated-measures ANOVA with Tukey’s HSD post-hoc tests and statistical significance defined as *p* < 0.05. Significant ANOVA effects are indicated in the upper left of each panel: Gas (*); Drug (#); Gas × drug interaction (†). Detailed statistics are reported in the Results, and post-hoc comparisons are listed in Table 2.

Naloxone hydrochloride alone did not alter V̇_I_, V_T_, or breathing frequency at rest or in response to HH. Subsequent buprenorphine reduced V̇_I_ during RA and HH via frequency depression (Fig. 5B and 5D). Pre-treatment with naloxone hydrochloride prevented buprenorphine-induced depression of V_T_ (Fig. 5C). This contrasts with Experiment I, where naloxone hydrochloride post-treatment failed to restore V_T_ to baseline during HH (Fig. 2C). *Respiratory pattern*

Two-way repeated measures ANOVA revealed a significant effect of drug (F_2,18_ = 23.17, *p* < 0.001) and a gas × drug interaction (F_2,18_ = 5.83, *p* = 0.0119) for T_I_, which was unaffected by naloxone hydrochloride alone but increased following buprenorphine (Fig. 6B). Significant effects of gas mixture (F_1,,9_ = 25.64, *p* < 0.001) and drug (F_2,18_ = 14.30, *p* > 0.001) were observed for T_E_. Naloxone hydrochloride alone did not affect T_I_ or T_E_ during RA breathing or HH. It also did not prevent buprenorphine-induced prolongation of T_I_ and T_E_, which was most pronounced during HH (Fig. 6C). Consistent with Experiment I, there were no significant effects of gas mixture or drug on apnea frequency or duration. Under control conditions, apnea frequency during RA breathing averaged 0.7 ± 0.4 apneas per minute, with a mean duration of 0.6 ± 0.3 s.

**Figure 6.**
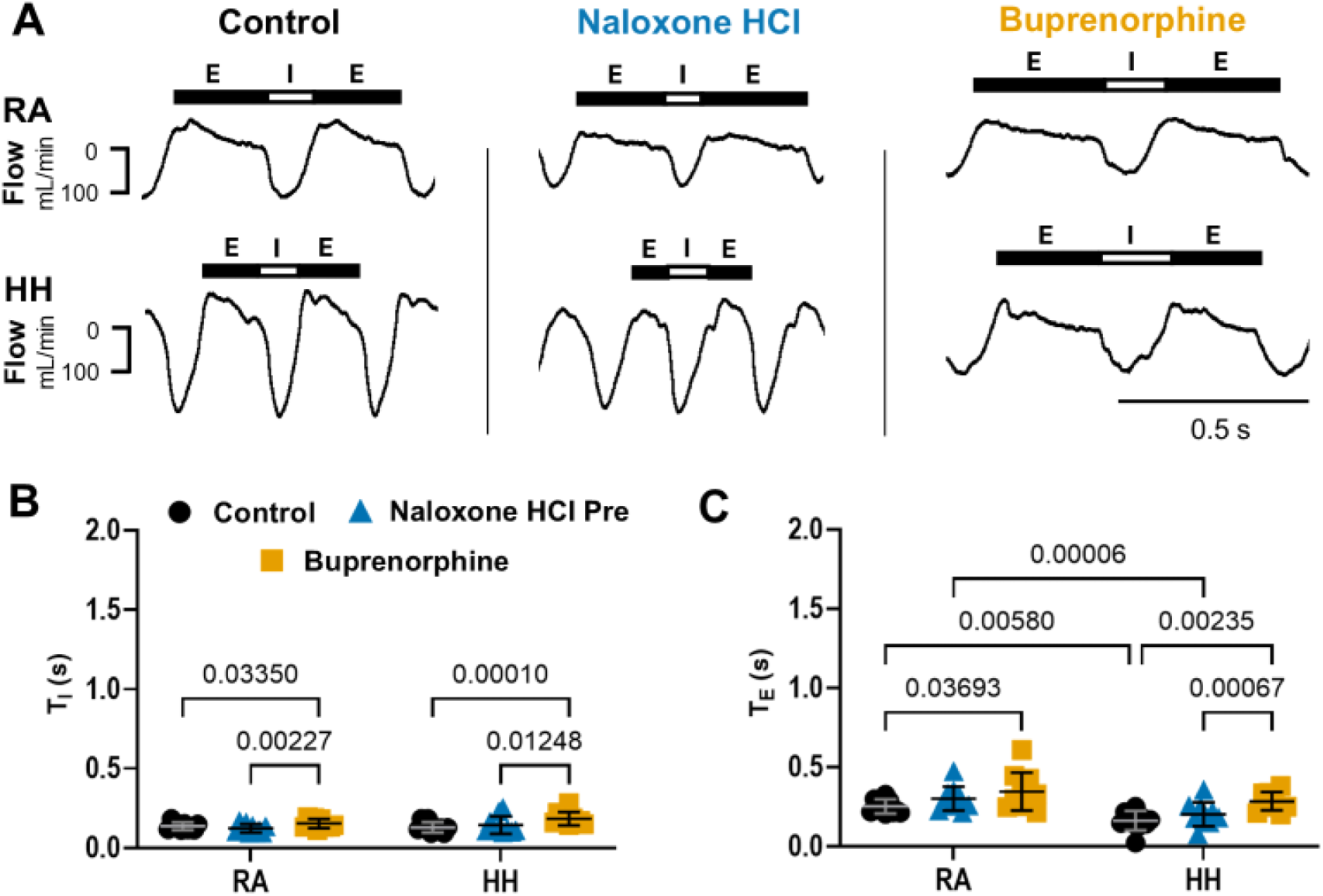
Naloxone hydrochloride pre-treatment does not prevent buprenorphine-induced breathing-phase prolongation. T_I_ and T_E_ were measured using dual-chamber plethysmography in neonatal Sprague Dawley rats (*n*=10, 5 male; P4-5) during RA breathing and HH. Measurements were obtained before and after naloxone hydrochloride administration, followed by buprenorphine. **Panel A:** Representative plethysmography traces illustrate breathing patterns across drug conditions (Control, Naloxone Hydrochloride, Buprenorphine) and gas mixtures (RA, HH). Troughs and plateaus represent inspiration (I) and expiration (E), respectively. **Panels B and C:** Plots show group data for T_I_ (B) and T_E_ (C). T_I_ and T_E_ were unaffected by naloxone hydrochloride alone but increased following subsequently administered buprenorphine, eliminating T_E_ shortening in response to HH. Data are presented as mean ± SD. Statistical analyses were performed using two-way repeated-measures ANOVA with Tukey’s HSD post-hoc tests and statistical significance defined as *p* < 0.05. Significant post-hoc comparisons are indicated on the plots.

#### Metabolic rate

One-way repeated measures ANOVA revealed a significant effect of drug for V̇O₂ (F_2,18_ = 3.49, *p* = 0.0463), V̇CO₂ (F_2,18_ = 4.29, *p* = 0.0316), V̇_I_/V̇O₂ (F_2,18_ = 6.92, *p* = 0.00648), and V̇_I_/V̇CO₂ (F_2,18_ = 10.90, *p* < 0.001). V̇O₂ and V̇CO₂ decreased following naloxone hydrochloride and were unchanged by subsequent buprenorphine. V̇I/V̇O₂ and V̇I/V̇CO₂ increased following naloxone hydrochloride alone, indicating hyperventilation. Naloxone hydrochloride pre-treatment prevented reductions in V̇I/V̇O₂ and V̇I/V̇CO₂ following subsequent buprenorphine administration. This contrasts with Experiment I, where buprenorphine reduced V̇I/V̇O₂ and V̇I/V̇CO₂, and naloxone hydrochloride post-treatment failed to restore V̇I/V̇O₂ and V̇I/V̇CO₂ to baseline levels. Data and significant post-hoc comparisons are shown in Figure 7.

**Figure 7.**
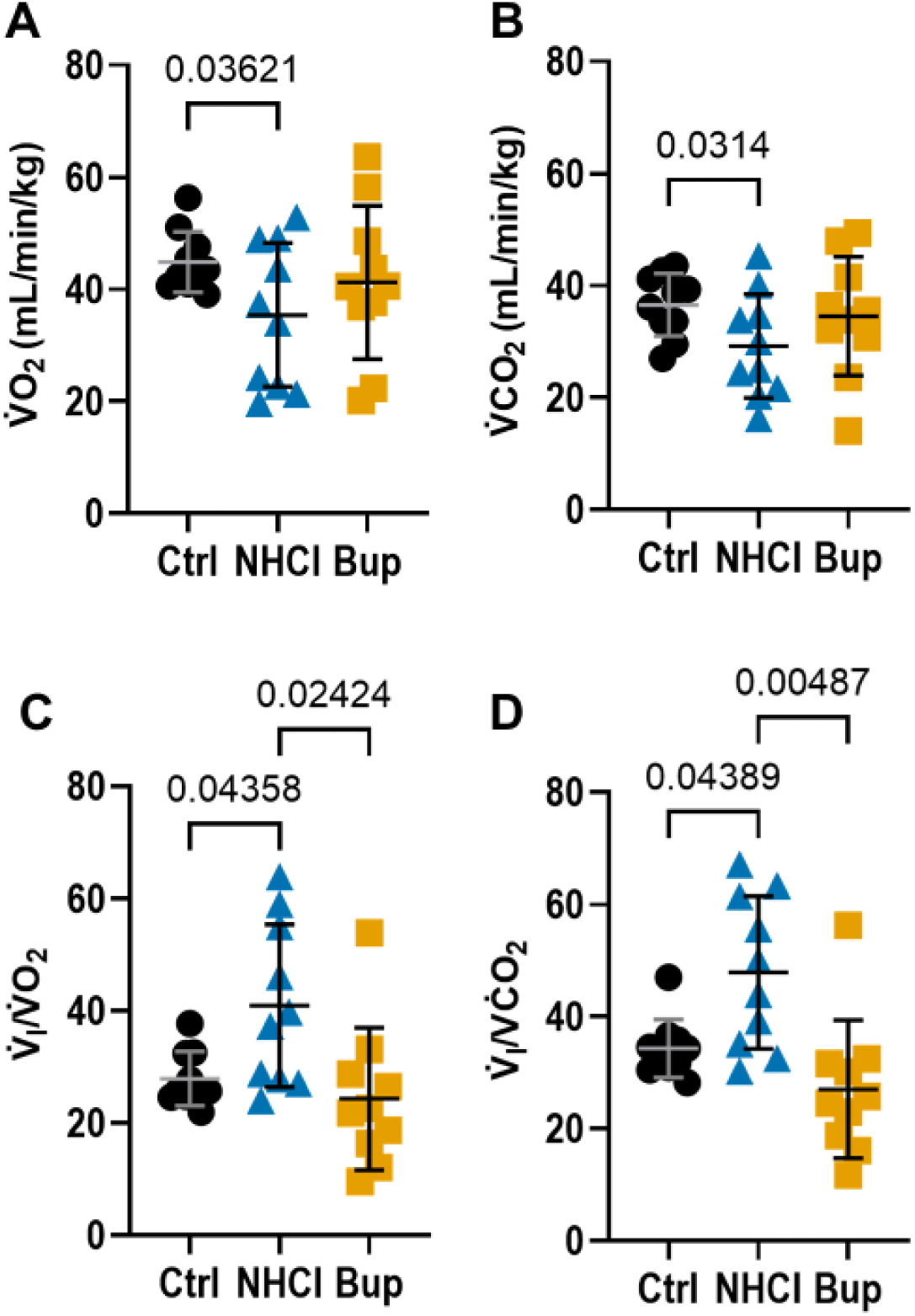
Naloxone hydrochloride alone reduces metabolic rate and increases ventilatory equivalents for O_2_ and CO_2_, indicating hyperventilation. V̇O₂, V̇CO₂, V̇I/V̇O₂ and V̇I/V̇CO₂ were measured using dual-chamber plethysmography in neonatal Sprague Dawley rats (*n*=10, 5 male; P4-5) during RA breathing. Measurements were obtained under control conditions (Ctrl) and repeated following administration of naloxone hydrochloride (NHCl), followed by buprenorphine (Bup). **Panels A-D:** Group data showing V̇O₂ (A), V̇CO₂ (B), V̇I/V̇O₂ (C) and V̇I/V̇CO₂ (D) across drug conditions. Naloxone alone reduced V̇O₂ and V̇CO₂, with no further change following subsequent buprenorphine administration. V̇_I_/V̇O_2_ and V̇_I_/V̇CO_2_ increased following naloxone alone, indicating hyperventilation. Notably, naloxone pre-treatment prevented the reduction of V̇_I_/V̇O_2_ and V̇_I_/V̇CO_2_ following subsequent buprenorphine administration. Data are presented as mean ± SD. Statistical analyses were performed using one-way repeated-measures ANOVA with Tukey’s HSD post-hoc tests and statistical significance defined as *p* < 0.05. Significant post-hoc comparisons are indicated on the plots.

### Experiment III: Naloxone methiodide followed by buprenorphine

#### V̇_I_, V_T_ and breathing frequency

Two-way repeated measures ANOVA revealed a significant effect of gas mixture for V̇_I_ (F_1,7_ = 305.9, *p* < 0.0001) and V_T_ (F_1,7_ = 185.1, *p* < 0.001). Significant effects of gas mixture (F_1,7_ = 67.98, *p* < 0.0001), drug (F_2,14_ = 45.56, p < 0.0001), and a gas × treatment interaction (F_2,14_ = 27.04, *p* < 0.0001) were observed for breathing frequency. Data are shown in Figure 8, with post-hoc comparisons listed in Table 3.

**Figure 8.**
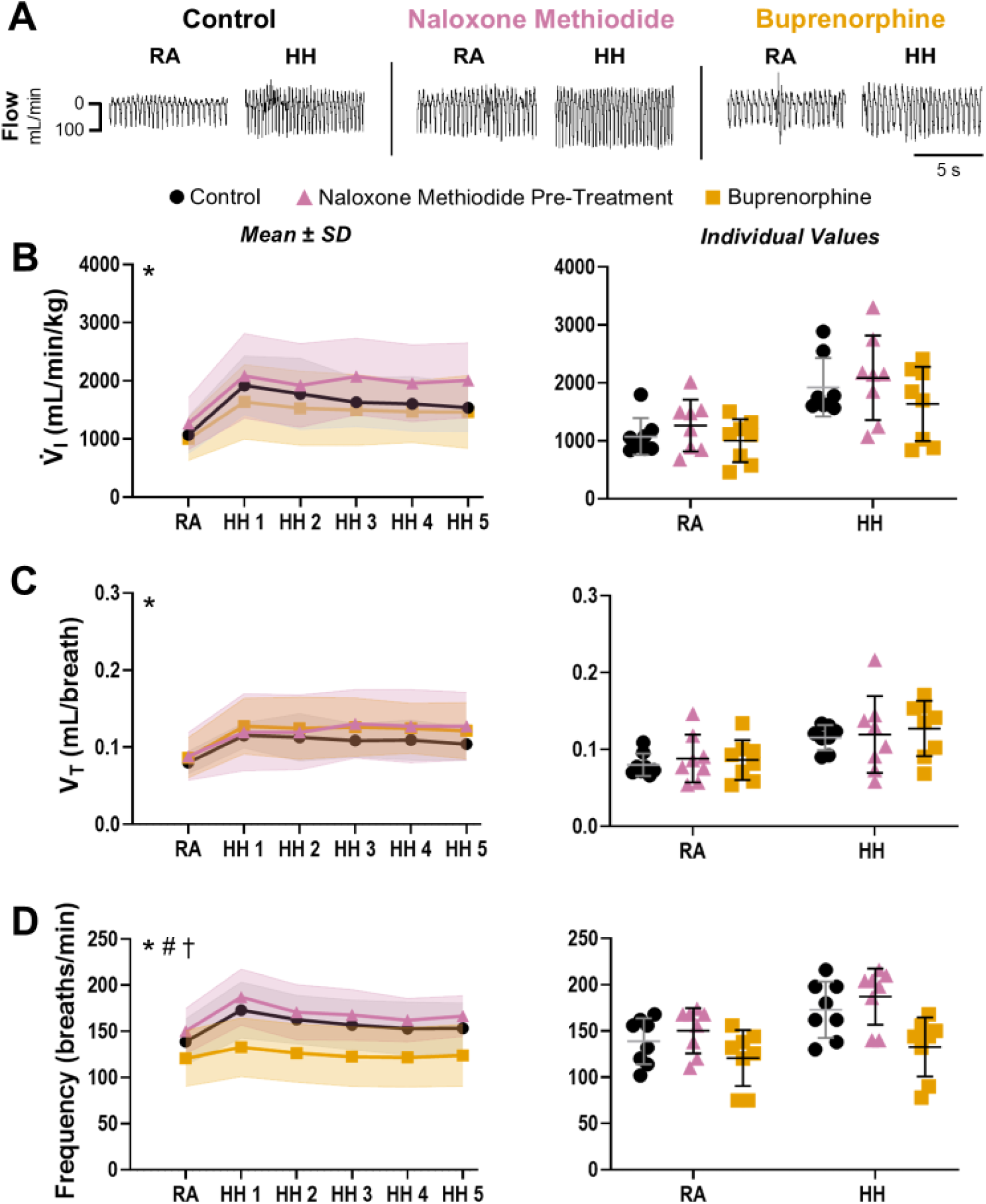
Naloxone methiodide pre-treatment mitigates buprenorphine-induced tidal volume depression. V̇_I_, V_T_ and breathing frequency were measured using dual-chamber plethysmography in neonatal Sprague Dawley rats ( *n*= 8, 4 male; P4-5) during RA breathing and a 5-min HH challenge (HH1-5). Measurements were obtained before and after administration of naloxone methiodide, followed by buprenorphine. **Panel A:** Representative plethysmography traces illustrate the effects of gas mixture and drug condition on breathing. **Panels B–D:** Group data (left) with individual values (right) show the effects of drug and gas mixture on V̇_I_ (B), V_T_ (C), and breathing frequency (D). Naloxone methiodide alone did not affect V̇_I_, V_T_, or breathing frequency during RA or HH. Buprenorphine subsequently reduced breathing frequency during both RA and HH. Naloxone methiodide pre-treatment prevented buprenorphine-induced depression of V_T_, resulting in V̇_I_ values that remained unchanged relative to control following buprenorphine. Data are presented as mean ± SD. Statistical analyses were performed using two-way repeated-measures ANOVA with Tukey’s HSD post-hoc tests and statistical significance defined as *p* < 0.05. Significant ANOVA effects are indicated in the upper left of each panel: Gas (*); Drug (#); Gas × drug interaction (†). Detailed statistics are reported in the Results, and post-hoc comparisons are listed in Table 3.

Naloxone methiodide alone did not alter V̇_I_, V_T_, or breathing frequency at rest or during the HH challenge (Fig. 8). Subsequent buprenorphine injection reduced breathing frequency during both RA and HH (Fig. 8D). Notably, naloxone methiodide pre-treatment mitigated buprenorphine-induced depression of V̇_I_ by preserving V_T_ (Fig. 8B and 8C), as did naloxone hydrochloride pre-treatment (Fig. 5C).

#### Respiratory pattern

Two-way repeated measures ANOVA revealed a significant effect of drug (F_2,14_ = 9.742, *p =* 0.00814) for T_I_, which was unaffected by naloxone methiodide and increased following buprenorphine administration (Fig. 9B). Significant effects of gas mixture (F_1,7_ = 66.15, *p* < 0.0001) and drug (F_2,14_ = 9.33, *p* = 0.0108) were observed for T_E_. Naloxone methiodide alone did not affect T_E_, and it did not prevent buprenorphine-induced T_E_, which persisted during the HH challenge (Fig. 9C). Consistent with Experiments I and II, there were no significant effects of gas mixture or drug on apnea frequency or duration. Under control conditions, apnea frequency during RA breathing averaged 0.8 ± 0.6 apneas per minute, with a mean duration of 0.8 ± 0.5 s.

**Figure 9.**
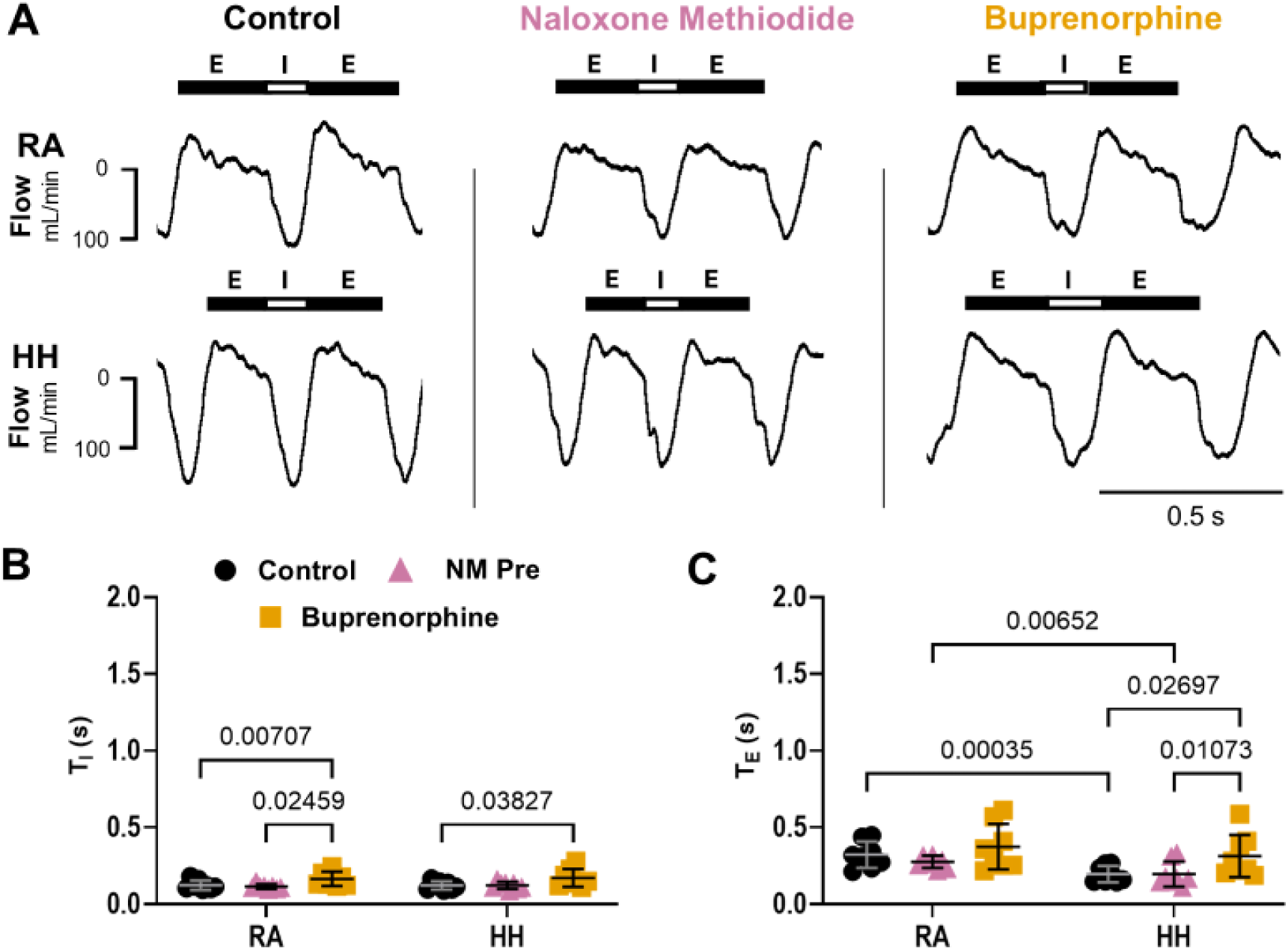
Naloxone methiodide pre-treatment does not prevent buprenorphine-induced breathing phase prolongation. T_I_ and T_E_ were measured using dual-chamber plethysmography in neonatal Sprague Dawley rats (*n* = 8, 4 male; P4-5) during RA breathing and HH. Measurements were obtained before and after naloxone methiodide administration, followed by buprenorphine. **Panel A:** Representative plethysmography traces illustrate breathing patterns across drug conditions Control, Naloxone Methiodide, Buprenorphine) and gas mixtures (RA, HH). Troughs and plateaus represent inspiration (I) and expiration (E), respectively. **Panels B and C:** Group data showing T_I_ (B) and T_E_ (C). T_I_ and T_E_ were unaffected by naloxone methiodide alone (NM Pre), but increased following subsequent buprenorphine administration, which abolished T_E_ shortening in response to HH. Data are presented as mean ± SD. Statistical analyses were performed using two-way repeated-measures ANOVA with Tukey’s HSD post-hoc tests and statistical significance defined as *p* < 0.05. Significant post-hoc comparisons are indicated on the plots.

#### Metabolic rate

Naloxone methiodide alone had no significant effect on V̇O₂, V̇CO₂, V̇_I_/V̇O₂, or V̇_I_/V̇CO₂. As with naloxone hydrochloride (Experiment II), naloxone methiodide prevented depression of V̇O₂ and V̇CO₂ following subsequent buprenorphine administration. Remarkably, V̇_I_/V̇O₂ and V̇_I_/V̇CO₂ were unchanged by naloxone methiodide alone, whereas naloxone hydrochloride induced hyperventilation. Naloxone methiodide pre-treatment prevented buprenorphine-induced reductions in V̇_I_/V̇O₂ and V̇_I_/V̇CO₂. Data and significant post-hoc comparisons are shown in Figure 10.

**Figure 10.**
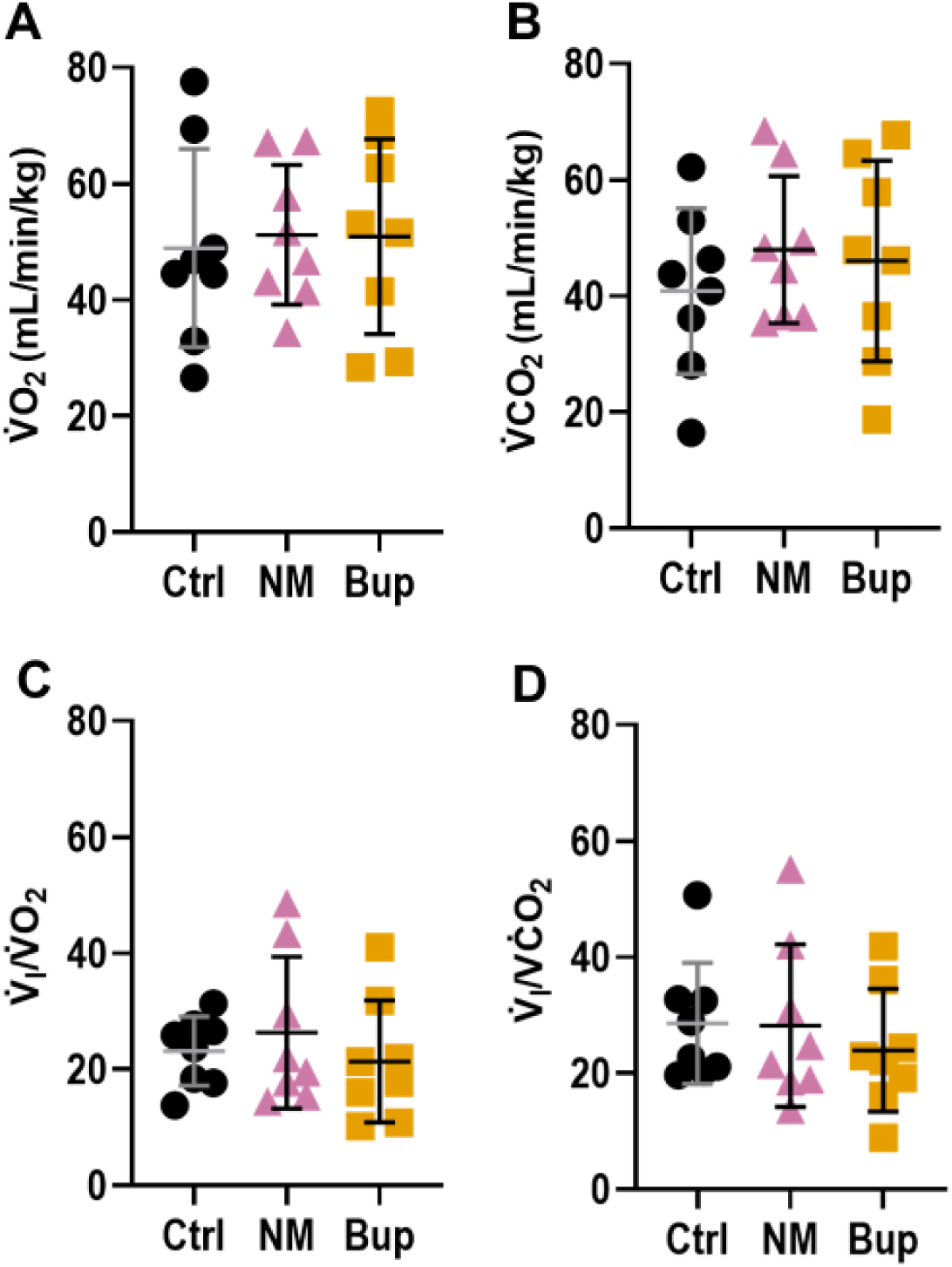
Naloxone methiodide pre-treatment does not affect metabolic rate, and prevents buprenorphine-induced hypoventilation. V̇O₂, V̇CO₂, V̇I/V̇O₂ and V̇I/V̇CO₂ were measured using dual-chamber plethysmography in neonatal Sprague Dawley rats (*n* = 8, 4 male; P4-5) during RA breathing. Measurements were obtained under control conditions (Ctrl) and repeated following administration of naloxone methiodide (NM), followed by buprenorphine (Bup). **Panels A-D:** Group data showing V̇O₂ (A), V̇CO₂ (B), V̇I/V̇O₂ (C) and V̇I/V̇CO₂ (D) across drug conditions. Naloxone methiodide pre-treatment did not alter V̇O₂ or V̇CO₂, and prevented their depression following subsequent buprenorphine administration. V̇_I_/V̇O_2_ and V̇_I_/V̇CO_2_ were likewise unaffected by naloxone methiodide pre-treatment and were not reduced following buprenorphine. This contrasts with Experiment II, where naloxone hydrochloride alone increased V̇_I_/V̇O_2_ and V̇_I_/V̇CO_2_, indicative of hyperventilation (Fig. 5). Data are presented as mean ± SD. Statistical analyses were performed using one-way repeated-measures ANOVA with Tukey’s HSD post-hoc tests and statistical significance defined as *p* < 0.05. Significant post-ho comparisons are indicated on the plots.

### Supplemental Analyses

Data from experiments in which naloxone methiodide was given after buprenorphine are included in the Supplemental Materials. The results are comparable to those of Experiment I, in that buprenorphine depressed V̇_I_ during RA and HH, and subsequent naloxone methiodide administration did not restore ventilatory parameters to baseline (Supplemental Figure 1, Supplemental Table 1). Time and sham-controls were conducted using two saline injections instead of the two drug injections. Two-way repeated measures ANOVA revealed no significant effect of saline injection, indicating that V̇_I_ was stable during RA and HH across repeated trials (Supplemental Figure 2). Finally, male and female neonatal rats (P4-5) exhibited comparable ventilatory responses to HH and buprenorphine (Supplemental Figure 3).

## Discussion

We found that acute buprenorphine exposure depresses V̇_I_ and metabolic rate during eupnea and profoundly blunts the HHVR in awake neonatal rats. Both V_T_ and breathing frequency were reduced following buprenorphine. Frequency depression was driven by prolonged T_I_ and T_E_. Once established, buprenorphine-induced respiratory depression was resistant to reversal by opioid receptor antagonists. In contrast, pretreatment with either naloxone hydrochloride or peripherally restricted naloxone methiodide mitigated buprenorphine-induced respiratory depression. Our results demonstrate that the developing respiratory control network is depressed by buprenorphine and suggests that peripheral opioid receptors contribute to buprenorphine-induced respiratory depression. Together, these findings provide novel insight into mechanisms of respiratory instability in neonates exposed to buprenorphine and suggest that pre-treatment with peripheral opioid receptor antagonists may offer a strategy to attenuate opioid induced respiratory depression (OIRD) without inducing withdrawal symptoms.

### Characteristics of buprenorphine-induced respiratory depression

The ability to increase V̇_I_ in response to reduced PaO_2_ and elevated PaCO_2_ is a critical homeostatic reflex across the lifespan (Darnall, 2010; Teppema & Dahan, 2010). This response is particularly important during sleep, when apneas and airway obstruction are more likely to occur (Cohen & Katz-Salamon, 2005). Notably, we found that buprenorphine significantly depressed the HHVR at a dose that did not cause apnea in eupnea or during HH. This suggests that following buprenorphine exposure, the respiratory system may appear stable during eupnea, but lacks the ability to compensate when challenged. Consistent with previous reports that both acute and chronic opioid exposure depress neonatal chemoreflexes (Colman & Miller, 2002; Berner *et al*., 2012; Osborne *et al*., 2022), our results suggest that impaired chemoreflex function may contribute to adverse neonatal outcomes following perinatal buprenorphine exposure (Kahila *et al*., 2007; Wallisch *et al*., 2010).

We found that prolonged T_E_ is the major driver of buprenorphine-induced breathing frequency depression, though reduced V_T_ also contributed to the depression of breathing following acute buprenorphine administration, both at rest and during HH. We also found that both V̇O_2_ and V̇CO_2_ decreased following buprenorphine administration. In the context of existing literature, we speculate that reduced respiratory muscle activity, together with bradycardia and decreased cardiac output, may limit tissue oxygen delivery and thereby reduce V̇O_2_ and V̇CO_2_ (Kesavan *et al*., 2014; Chen & Ashburn, 2015; Duursma *et al*., 2025). Opioids may also exert direct depression of cellular oxidative phosphorylation, reducing ATP levels and mitochondrial oxygen consumption (Tarazi & Maynes, 2023).

Importantly, the magnitude of respiratory depression exceeded the reduction in metabolic rate, as evidenced by decreased V̇I/V̇O₂ and V̇I/V̇CO₂. Together, these results indicate that in addition to profound chemoreflex depression, buprenorphine produces significant hypoventilation at rest, which could compromise oxygen delivery to the tissues even in the absence of frank apneic episodes. These observations are consistent with data in human subjects showing that buprenorphine is associated with sleep-disordered breathing and hypoxemia in adults (Farney *et al*., 2013), and reduced fetal breathing movements (Bulger *et al*., 2025). Together, these findings underscore the need to carefully consider the respiratory and cardiovascular consequences of buprenorphine administration, particularly in vulnerable or medically fragile populations.

### Effects of Naloxone Hydrochloride

Naloxone hydrochloride (Narcan) is a competitive opioid receptor antagonist that freely crosses the blood-brain barrier (Berkowitz, 1976). In Experiment I, naloxone hydrochloride was administered following buprenorphine to determine whether buprenorphine-induced hypoventilation could be reversed, reflecting the standard clinical practice of naloxone administration as a rescue therapy for opioid overdose. During RA breathing, naloxone hydrochloride post-treatment produced small but significant increases in V_T_ and breathing frequency, but these effects were insufficient to restore V̇_I_ to baseline. During the HH challenge, naloxone hydrochloride increased breathing frequency relative to buprenorphine alone, but frequency remained markedly below control levels. Moreover, naloxone hydrochloride post-treatment failed to increase V_T_ relative to buprenorphine during HH. V̇I/V̇O₂ and V̇I/V̇CO₂ likewise remained depressed following naloxone hydrochloride post-treatment. Overall, these results indicate that buprenorphine-induced hypoventilation and chemoreflex depression are resistant to reversal by naloxone hydrochloride. Despite its classification as a partial agonist, buprenorphine exhibits ultra-high affinity and slow dissociation kinetics at the *μ*-opioid receptor, (Volpe *et al*., 2011; Blazes & Morrow, 2020) suggesting that once bound, these properties prevent its competitive displacement by naloxone.

Naloxone hydrochloride alone (Experiment II) did not significantly change V̇_I_, V_T_, frequency, or the HHVR, suggesting that endogenous opioids do not influence these specific parameters in neonatal rats under the conditions examined. However, naloxone hydrochloride alone did reduce V̇O_2_ and V̇CO_2_, while increasing both V̇I/V̇O₂ and V̇I/V̇CO₂, consistent with hyperventilation. Previous investigations into the influence of endogenous opioids on ventilation and metabolic rate have yielded mixed results, reflecting methodological variability as well as differences in species, developmental stage, and the inherent complexity of the endogenous opioid system. (Malin *et al*., 1985; Kraus *et al*., 1996; Schlenker *et al*., 1997; Whitaker-Fornek & Levitt, 2025) The mechanisms underlying the reduction in V̇O_2_ and V̇CO_2_ and the relative hyperventilation observed following naloxone hydrochloride administration in the present study remain unclear and warrant further investigation.

Given that naloxone hydrochloride was minimally effective in reversing the effects of buprenorphine, it is interesting that pre-treatment in Experiment II significantly attenuated the magnitude of buprenorphine-induced hypoventilation by mitigating depression of V_T_. In other words, naloxone hydrochloride is more effective at preventing buprenorphine-induced hypoventilation than reversing it. The observation that buprenorphine still exerted a partial effect may reflect competitive displacement of naloxone hydrochloride at specific sites, or a fraction of receptors that remained available for buprenorphine binding.

### Peripheral mechanisms of opioid induced respiratory depression (OIRD)

Opioid receptors are a diverse family of inhibitory G protein-coupled receptors (GPCRs) expressed throughout the central and peripheral nervous system (Standifer & Pasternak, 1997; Stein & Lang, 2009). OIRD is generally attributed to activation of μ-opioid receptors located within the brainstem respiratory network, including central chemosensory areas (Pokorski *et al*., 1981; Hurle *et al*., 1985; Ballanyi *et al*., 1997; Lalley, 2003; Lorier *et al*., 2010; Phillips *et al*., 2012). Compared to central mechanisms, the contribution of peripheral opioid receptors to OIRD is less well understood. It is hypothesized that activation of opioid receptors on oxygen-sensing glomus cells in the carotid bodies reduces their excitability, thereby attenuating the hypoxic ventilatory response (Spiller *et al*., 2026). Anatomical evidence indicates that glomus cells express μ-, κ-, and δ-opioid receptors in neonatal rats, and calcium imaging shows that the μ-opioid receptor agonist DAMGO inhibits voltage-gated influx of calcium into these cells (Ricker *et al*., 2015). These and other observations showing evidence for a functional role of carotid body opioid receptors has generated interest in targeting peripheral opioid receptors to counter OIRD (McQueen & Ribeiro, 1980; Pokorski & Lahiri, 1981; Kirby & McQueen, 1986; Peng *et al*., 2025).

Quaternary naloxone methiodide is thought not to cross the blood brain barrier and has been shown to reverse opioid-induced brain hypoxia and cardiorespiratory depression without entering brain tissue (Perekopskiy *et al*., 2020; Ruyle *et al*., 2025) or inducing withdrawal (Lewanowitsch & Irvine, 2002). We found that naloxone methiodide pre-treatment prevented buprenorphine-induced depression of V_T_, resulting in preservation of V̇_I_ at rest and during HH. The primary protective effects of both naloxone methiodide and naloxone hydrochloride were on V_T_, suggesting that V_T_ depression by buprenorphine is mediated, at least in part, by peripheral opioid receptors. This conclusion is supported by prior evidence indicating that hypoxic stimulation of peripheral chemoreceptors increases V_T_ and that V_T_ is a major driver of the hypoxic ventilatory response in neonatal rats (Chiocchio *et al*., 1984; Fung *et al*., 1996; Zhao *et al*., 2016; Frazure *et al*., 2026). Naloxone methiodide pre-treatment also prevented buprenorphine-induced metabolic depression. Because naloxone methiodide alone did not affect V̇_I_/V̇O₂ or V̇_I_/V̇CO₂, we suggest that by mitigating respiratory depression (and the associated reduction in the work of breathing), naloxone methiodide prevented the decline in V̇O_2_ and V̇CO_2_ that is typically observed with buprenorphine.

Overall, our findings add to evidence supporting the utility of peripherally restricted antagonists for mitigating OIRD and extend this work by evaluating the efficacy of naloxone methiodide in countering buprenorphine-induced hypoventilation in neonatal rats. We show that although buprenorphine’s ultra-high binding affinity limits reversal once established (Supplemental Fig. 1), pre-treatment with naloxone methiodide markedly reduced the severity of respiratory depression following buprenorphine administration. These results have important implications for the development of strategies to stabilize breathing in medically fragile populations, as well as in patients receiving long-term opioid therapy.

### Sex and age as biological variables

Sex-specific effects of opioids are complex and remain poorly understood (Sarton *et al*., 2000; Cepeda & Carr, 2003; Cepeda *et al*., 2003; Bijur *et al*., 2008; Fullerton *et al*., 2018). In humans, morphine has been shown to reduce peripheral chemoreflex responses more significantly in women than in men (Sarton *et al*., 1999). Recent preclinical studies have shown that adult female rats exhibited greater respiratory depression following heroin (Marchette *et al*., 2023) and buprenorphine (Frazure *et al*., 2024) than males. In the present study, there were no sex-specific effects of buprenorphine on breathing in P4-5 neonatal rats (Supplemental Figure 3). The sustained apnea and mortality previously reported in adult female, but not male, rats following buprenorphine administration (Frazure *et al*., 2024) may arise from interactions between opioids and age-dependent hormonal influences that have not yet emerged by P4-5. More broadly, these findings highlight the need to consider drug type, developmental stage, species, and experimental setting when evaluating sex differences.

### Critique of the methods

Here, we asked whether buprenorphine impairs a neonate’s ability to respond appropriately to heightened respiratory drive. This question was addressed using a combined HH challenge. The answer was unambiguous, as buprenorphine produced a profound depression of the HHVR. Future studies could employ isolated hypoxic and hypercapnic challenges to distinguish peripheral vs. central mechanisms of opioid-induced chemoreflex depression.

The extent of naloxone methiodide’s peripheral selectivity remains controversial (Brown & Goldberg, 1985). Large systemic doses have been found to penetrate the blood-brain barrier to a limited extent, with brain concentrations ∼40-fold lower than those of naloxone hydrochloride (Perekopskiy *et al*., 2020). Accordingly, because blood and plasma concentrations of naloxone methiodide were not directly measured in the present study, a small degree of central penetration cannot be entirely ruled out.

Nevertheless, naloxone methiodide reverses opioid-induced respiratory depression in adult rats without detectable central penetration at doses of 1 or 5 mg/kg (Ruyle *et al*., 2025), and the dose used in the present study was at the low end of that range. Although the blood-brain barrier is still developing during the neonatal period, it is functionally protective from embryonic stages and undergoes substantial maturation by postnatal days 4-5 (Schulze & Firth, 1992; Saunders *et al*., 2012). Therefore, while the effects of naloxone methiodide were consistent with selective antagonism of peripheral opioid receptors and differed from those of naloxone hydrochloride, the results of Experiments III and IV should be interpreted with the caveat that blood-brain barrier permeability to naloxone methiodide was not directly assessed.

Our experiments were conducted in healthy, opioid-naïve neonatal rats. We acknowledge that responses to acute opioid receptor agonists and antagonists may differ in pathological contexts. Having established a foundational understanding of buprenorphine’s effects on neonatal respiratory control in normal development, it will be important to extend this work to include models that incorporate prenatal opioid exposure, postnatal opioid treatment, and comorbid inflammatory or injury-related conditions.

### Clinical relevance

Many studies devoted to understanding the respiratory effects of opioids have focused on mechanisms of opioid overdose. We advocate that the respiratory depressant effects of opioids in clinical settings are also important to understand, particularly in vulnerable or medically fragile populations. The present findings demonstrate that buprenorphine induces hypoventilation and attenuates chemoreflex function in neonates in the absence of overdose symptoms such as an increased number and duration of apneic episodes, raising the possibility that these effects may contribute to adverse outcomes in buprenorphine-exposed infants, although further investigation is needed.

Strategies are needed to counter OIRD in the acute inpatient setting and in patients receiving long-term opioid therapy. While naloxone hydrochloride (Narcan) effectively reverses life-threatening overdose, it is unsuitable adjuvant therapy for these populations because it readily crosses the blood-brain barrier and could therefore induce opioid withdrawal.

Importantly, the finding that pre-treatment with the peripherally restricted opioid receptor antagonist naloxone methiodide attenuated buprenorphine-induced respiratory depression suggests a potential strategy to stabilize breathing while minimizing the risk of precipitated opioid withdrawal. This result complements prior work showing that pre-treatment with the partial 5-HT_1A_ agonist buspirone similarly attenuates buprenorphine-induced respiratory depression in adult rats (Frazure *et al*., 2024). Collectively, these findings may inform the development of interventions aimed at protecting respiratory function in patients treated with opioids who are at elevated risk for respiratory insufficiency, including opioid-exposed infants.

## Additional Information

### Data availability statement

All relevant data, associated protocols and materials are described within the published article and are available from the corresponding author upon reasonable request.

### Author contributions

All persons designated as authors qualify for authorship, and all those who qualify for authorship are listed.

M.F. and R. F. were responsible for conception and design of the work. M.F., K.P., E.S., E.F., and R. F. were responsible for acquisition, analysis and interpretation of data for the work. M.F., K.P., E.S., E.F., and R. F. were responsible for drafting the work or revising it critically for important intellectual content; approved the final version of the manuscript; and agree to be accountable for all aspects of the work in ensuring that questions related to the accuracy or integrity of any part of the work are appropriately investigated and resolved.

