## Supplemental Materials for "Acute buprenorphine exposure depresses neonatal respiratory chemoreflexes in the presence or absence of naloxone"

Supplemental Figures: 3
Supplemental Tables: 1

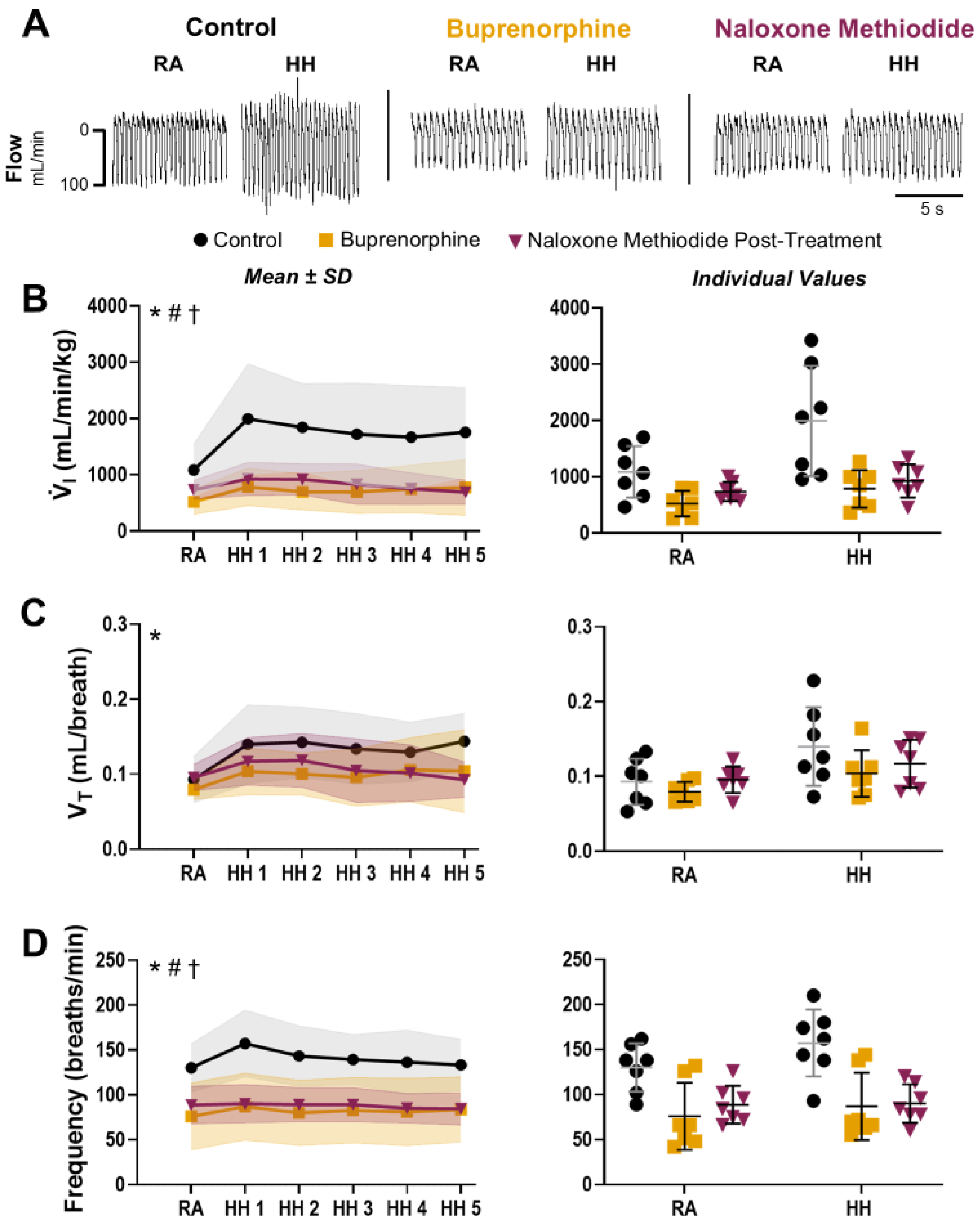

**Supplemental Figure 1. Naloxone methiodide post-treatment does not reverse buprenorphine-induced respiratory depression.** $\dot{V}_I$, $V_T$ and breathing frequency were measured using dual-chamber plethysmography in neonatal Sprague Dawley rats (*n* = 7, 4 male; P4-5) during RA breathing and a 5-min HH challenge (HH1-5). Measurements were obtained before and after administration of buprenorphine, followed by peripherally restricted naloxone methiodide. **Panel A:** Representative plethysmography traces illustrate the effects of gas mixture (RA, HH) and drug condition (Control, Buprenorphine, Naloxone Methiodide) on breathing. **Panels B–D:** Group data (left) and individual replicates (right) show the effects of drug and gas mixture on $\dot{V}_I$ (B), $V_T$ (C), and breathing frequency (D). Two-way repeated measures ANOVA revealed significant effects of gas ($F_{1,6} = 20.18$, $p = 0.00414$), drug ($F_{2,12} = 12.99$, $p = 0.00675$), and a gas × drug interaction ($F_{2,12} = 13.30$, $p = 0.00432$) for $\dot{V}_I$. A significant effect of gas mixture ($F_{1,6} = 12.98$, $p = 0.0113$) was observed for $V_T$. Significant effects of gas ($F_{1,6} = 22.11$, $p = 0.00332$), drug ($F_{2,12} = 31.33$, $p < 0.0001$), and gas × drug interaction ($F_{2,12} = 7.89$, $p = 0.00922$) were observed for breathing frequency. Buprenorphine reduced resting ventilation and markedly attenuated the ventilatory response to HH via blunted $V_T$ and elimination of the frequency response. Naloxone methiodide failed to restore $\dot{V}_I$ to baseline levels during RA or HH. Naloxone methiodide did not increase $V_T$ or breathing frequency relative to buprenorphine during HH. Data are presented as mean ± SD. Statistical analyses were performed using two-way repeated-measures ANOVA with Tukey's HSD post-hoc tests and statistical significance defined as $p < 0.05$. Significant ANOVA results are indicated in the upper left corner of each panel: Gas mixture (*); drug (#); and gas × drug interaction (†). Post-hoc comparisons are listed in Supplemental Table 1.

**Supplemental Table 1. Post-hoc comparisons of respiratory parameters for experiments where buprenorphine was followed by naloxone methiodide.** *P*-values shown in bold when significant ($p < 0.05$). *RA = Room air; HH = Hypoxic-hypercapnic challenge; Ctrl = Control; Bup = Buprenorphine; NM Post = Naloxone methiodide post-treatment.*

| **Gas** | **Drug Comparison** | **$\dot{V}_I$** | **$V_T$** | **Frequency** |
|---|---|---|---|---|
| | | *p-value* | *p-value* | *p-value* |
| RA | Ctrl vs. Bup | **0.0101** | 0.400 | **0.00120** |
| | Bup vs. NM Post | 0.0518 | **0.0227** | 0.359 |
| | NM Post vs. Ctrl | 0.129 | 0.967 | **0.00193** |
| HH | Ctrl vs. Bup | **0.00942** | **0.0467** | **0.00173** |
| | Bup vs. NM Post | 0.563 | 0.556 | 0.931 |
| | NM Post vs. Ctrl | **0.0391** | 0.411 | **0.00247** |
| **Drug** | **Gas Comparison** | **$\dot{V}_I$** | **$V_T$** | **Frequency** |
| | | *p-value* | *p-value* | *p-value* |
| Ctrl | RA vs. HH | **0.00508** | **0.00546** | **0.00554** |
| Bup | RA vs. HH | **0.0299** | **0.0438** | 0.0614 |
| NM Post | RA vs. HH | 0.0593 | 0.0665 | 0.617 |

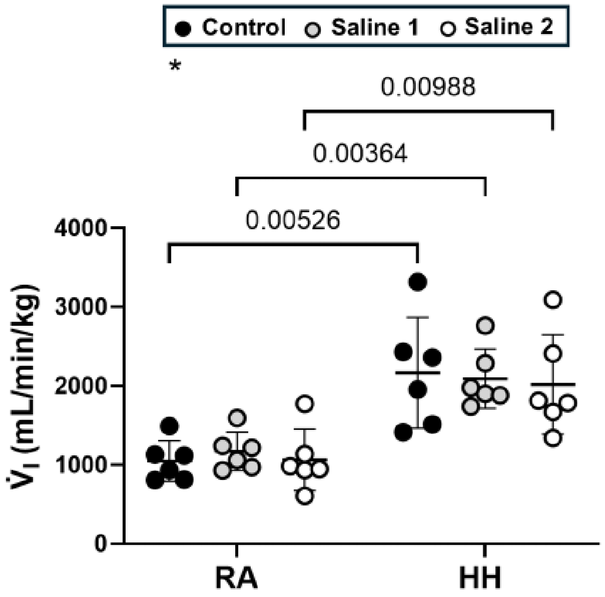

**Supplemental Figure 2. Ventilation is unaffected by sham injections and remains stable across repeated trials.** In time and sham-controls, $\dot{V}_I$ was measured using dual-chamber plethysmography in neonatal Sprague Dawley rats (n = 6; 3 males; P4-5) during RA breathing and a HH challenge. Measurements were obtained at baseline, and after each of two sham injections (Saline 1, Saline 2) over a time course matched to Experiments I-III. Data are shown as individual replicates with mean ± SD for each gas and treatment condition. Two-way repeated measures ANOVA revealed a significant effect of gas mixture ($F_{1,10} = 45.19$, $p = 0.00110$), but no effect of saline injection. $\dot{V}_I$ consistently increased in response to HH, with no difference between trials. Significant ANOVA results are indicated in the upper left corner: Main effect of gas mixture (*). Significant post-hoc comparisons (Tukey's HSD) are shown on the plot. The threshold for statistical significance was $p < 0.05$.

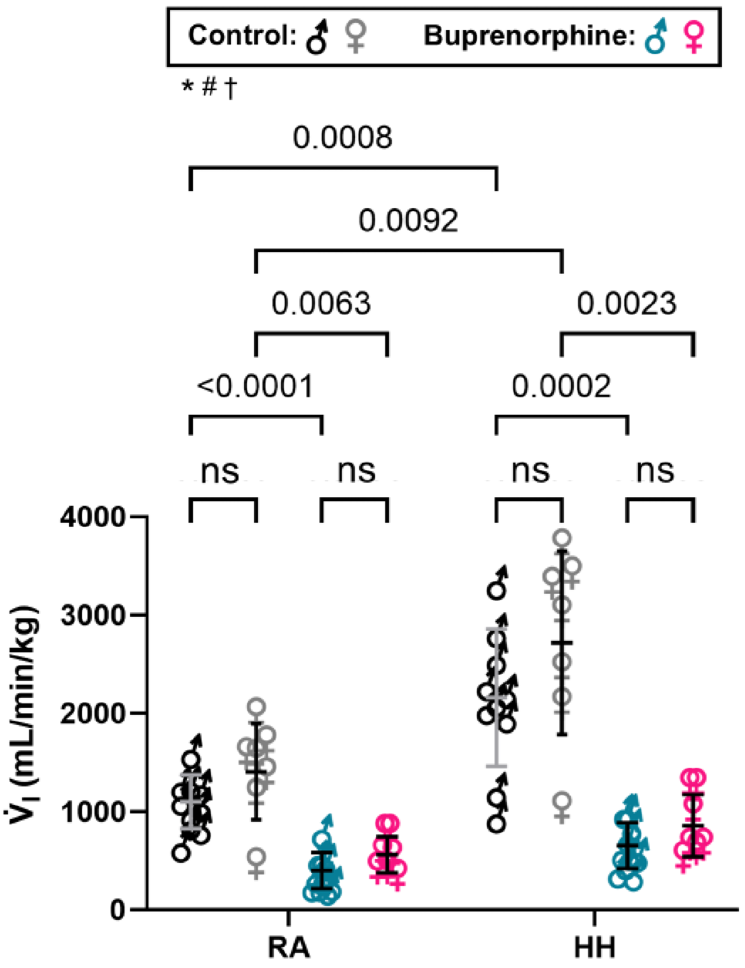

**Supplemental Figure 3. Acute buprenorphine depresses the neonatal hypoxic-hypercapnic ventilatory response independent of sex.** $\dot{V}_I$ was measured in male ($n$ = 10) and female ($n$ = 7) P4–5 Sprague Dawley rat pups during RA breathing and a HH challenge at baseline and following buprenorphine administration. Data are shown as individual values with mean ± SD for both sexes under each gas and treatment condition. A three-way repeated-measures ANOVA revealed significant effects of gas mixture ($F_{1,15}$ = 88.89, $p$ < 0.0001), drug ($F_{1,15}$ = 111.4, $p$ < 0.0001), and a gas × drug interaction ($F_{1,15}$ = 53.89, $p$ < 0.0001). There was no significant effect of biological sex. At baseline, $\dot{V}_I$ increased significantly in response to HH in both males and females. Following buprenorphine, $\dot{V}_I$ was reduced during RA and HH in both males and females. Significant ANOVA effects are indicated in the upper left corner: Gas mixture (*); drug (#); gas × drug interaction (†). Significant post-hoc comparisons (Tukey's HSD) are shown on the plot. The threshold for statistical significance was $p < 0.05$.
